# The wheat NLR protein PM3b localizes to endoplasmic reticulum-plasma membrane contact sites and interacts with AVRPM3^b2/c2^ through its LRR domain

**DOI:** 10.1101/2024.09.27.615346

**Authors:** Jonatan Isaksson, Lukas Kunz, Simon Flückiger, Victoria Widrig, Beat Keller

**Affiliations:** Department of Plant and Microbial Biology, University of Zurich, Zollikerstrasse 107, 8008 Zurich, Switzerland; Athebio AG, Grabenstrasse 11a, 8952 Schlieren, Switzerland; Department of Microbiology and Genetics, Spanish-Portuguese Agricultural Research Center (CIALE), University of Salamanca, 37007 Salamanca, Spain

**Keywords:** plant immunity, ER-PM contact sites, pathogen effector, powdery mildew, wheat, plant-fungal interactions, NLR

## Abstract

Plant nucleotide-binding leucine-rich repeat (NLR) proteins are intracellular immune receptors that directly or indirectly perceive pathogen derived effector proteins to induce an immune response. NLRs display diverse sub-cellular localizations, which are associated with the capacity of the immune receptor to confer disease resistance and recognize its corresponding avirulence effector. In wheat, the NLR PM3b recognizes the wheat powdery mildew effector AVRPM3^b2/c2^ and we examined the molecular mechanism underlying this recognition. We show that PM3b and other PM3 variants localize to endoplasmic reticulum (ER)-plasma membrane (PM) contact sites (EPCS) while AVRPM3^b2/c2^ localizes to the nucleocytoplasmic space. Additionally, we found that PM3b interacts in planta with AVRPM3^b2/c2^ through its LRR domain. We further demonstrate that full length PM3b interaction with AVRPM3^b2/c2^ is considerably weaker than for the isolated PM3b LRR domain or the susceptible PM3 variant PM3CS, indicating that activation of PM3b leads to dissociation of the complex. In line with this: We observed a strong interaction between PM3b and AVRPM3^b2/c2^ in a P-loop mutant of PM3b which was unable to initiate a cell death response, or when an inactive variant of AVRPM3^b2/c2^ was used. We propose that PM3b transiently interacts with AVRPM3^b2/c2^ through residues in the LRR which are conserved among PM3 variants while the amino acids necessary for full activation and cell death signaling are unique to PM3b. Our data suggests that PM3b localization and interaction with AVRPM3^b2/c2^ differs from other well studied NLRs and further highlights the mechanistic diversity in NLR-mediated responses against pathogens in plants.

## Introduction

Plants are protected from colonization by most microbes by immune receptors at the cell-surface which can detect broadly conserved molecular patterns, such as chitin in the fungal cell wall, resulting in pattern-triggered immunity (PTI) (DeFalco & Zipfel, 2021). Pathogenic fungi secrete hundreds of small proteins called effectors which can manipulate host protein machinery to suppress innate immune responses and divert cellular processes to invade their plant host (Ngou et al., 2022). To protect themselves from pathogens, plants have evolved intracellular immune receptors that guard the cell for potential intruders. Many of these immune receptors are nucleotide-binding leucine-rich repeat (NLR) proteins which directly or indirectly detect effectors, then termed avirulence (AVR) proteins, and induce effector-triggered immunity (ETI) (Chen et al., 2022). In contrast to bacteria, whose effectors have often been described to be indirectly recognized, direct effectors recognition is most often found for filamentous fungi (Chen et al., 2022).

Interestingly, while culminating in the same typical hypersensitive response (HR) and cell death phenotypes, the localization and its relationship to signaling function of NLR-type immune receptors is highly diverse. The NLR protein MLA10 plays a dual role in the immune response to barley powdery mildew by interacting with a transcription factor in the nucleus, while cytoplasmic localization is required for its ability to trigger cell death upon AVR recognition (Shen et al., 2007). Another NLR protein RPM1 localizes to the plasma membrane where it senses modifications to RIN4 by AVRRPM1 and initiates a cell death response (Gao et al., 2011). The NLR required for cell death (NRC) members NRC2 and NRC4 localize to cytoplasmic filaments or the cytoplasm, respectively, in their resting state but form puncta at the plasma membrane (PM) upon effector recognition (Contreras et al., 2023; Duggan et al., 2021).

The *PM3* allelic series in wheat consists of 17 functional alleles encoding NLRs with an N-terminal coiled-coil (CC) domain, which provide race-specific resistance against the wheat powdery mildew pathogen *Blumeria graminis* f. sp. *tritici* (*Bgt*). The variants encoded by the *Pm3* alleles show a high level of similarity among each other on the amino acid sequence level (>97 %) (Bhullar et al., 2009). Even so, *Pm3* alleles have been shown to be highly specific in their recognition of their corresponding *AvrPm3*s (Bourras et al., 2015, 2019). To date, *AvrPm3^a2/f2^*, *AvrPm3^b2/c2^* and *AvrPm3^d3^*have been cloned and are recognized by PM3a/PM3f, PM3b/PM3c and PM3d, respectively, as indicated by the induction of HR upon co-expression in *Nicotiana benthamiana*. In addition to the identified *AvrPm3* effector genes, *SvrPm3^a1/f1^* was identified which can suppress resistance mediated by all tested *Pm3* alleles (Bourras et al., 2016; Parlange et al., 2015). Interestingly, the sequence homology between the three AVRPM3 effector proteins is low but they are hypothesized to have a shared RNase-like structure, which might be why they are recognized by highly similar variants of PM3 (Bourras et al., 2019).

Similar observations have been made for the allelic series of barley resistance genes encoding the NLR-type immune receptor MLA. MLA variants are highly similar to each other but recognize sequence diverse avirulence effectors (AVRa) from the barley powdery mildew pathogen *Blumeria hordei* (*Bh*) (Saur et al., 2019). Crystal structures of AVR_A6,_ AVR_A7_-1, AVR_A10_ and AVR_A22_, recognized by MLA6, MLA7, MLA10 and MLA22, respectively, and the *Bgt* effector AVRPM2 which is recognized by the wheat immune receptor PM2 confirmed their predicted RNase-like structure (Cao et al., 2023). Furthermore, a direct interaction between MLA7, MLA10, MLA13, and MLA22 and AVR_a7_, AVR_a10_, AVR_a13,_ and AVR_a22_ was suggested based on Yeast-2 Hybrid and Split-Luciferase assays (Saur et al., 2019). This was recently confirmed with a cryo-EM structure of a MLA13-AVR_a13_-1 heterodimeric complex which showed that AVR_a13_-1 also has a RNase-like structure and makes multiple contacts with the LRR and the Winged Helix Domain (WHD) of MLA13 (Lawson et al., 2024). While much has been learned about the molecular mechanism of other mildew immune receptors such as MLAs and their corresponding AVRa effectors from *Bh*, much less is known about these mechanisms for PM3s and AVRPM3 effectors. The study by Cao and colleagues (2023) for example failed to produce crystals for the tested AVRPM3 effectors, although they are also likely to share the RNAse-like structure as indicated by structural modelling (Bourras et al., 2019). Furthermore, previous studies have revealed a complex genetic model for the relations between *Bgt* effector components and PM3 variants (Bourras et al., 2016) and our study aims to investigate the molecular mechanism of PM3 mediated defense responses in wheat against *Bgt*.

Here we describe the spatial distribution of PM3b and other PM3 variants which localize to specialized membrane contact sites at the endoplasmic reticulum (ER) network and plasma membrane (PM). We show that AVRPM3^b2/c2^ interacts with PM3b in planta through its LRR domain and that amino acids conserved between different PM3 variants play a role in this interaction which might explain why very few amino acid polymorphisms are needed between PM3 variants to recognize different AVRPM3 effectors.

## Results

### PM3b localizes into distinct puncta at the periphery of the ER-network and the PM

As a first step in understanding the molecular mechanism of PM3b mediated resistance in wheat, we wanted to investigate its subcellular localization in its resting state. To do this, we fused eGFP to the C-terminus of PM3b and transiently expressed it in *N. benthamiana* and wheat. Fusion of a large fluorescent tag to a NLR might lead to a non-functional protein. However, PM3b-eGFP could still trigger a hypersensitive response in *N. benthamiana* upon co-expression with the avirulent variant I of AVRPM3^b2/c2^ (hereafter AVRPM3^b2/c2^-I). Co-expression of an eGFP-tagged version of the susceptible variant PM3CS (Yahiaoui et al. 2006) with AVRPM3^b2/c2^-I did not elicit a cell death response and neither did AVRPM3^b2/c2^-I or PM3b alone (Supplementary Fig. 1a). Laser scanning confocal microscopy (LSCM) revealed that in its resting state PM3b formed distinct puncta throughout the cell that were not suggestive of localization to any of the major sub-cellular compartments. Co-expression of PM3b-eGFP with the ER marker SP-mRFP-HDEL (Nelson et al., 2007) revealed that the puncta partially co-localized with the ER network. Within the ER network, PM3b-eGFP mainly accumulated at three-way junctions and cisternae, and weakly to the tubules. (Fig. 1a). A line measurement of the brightness intensity of the pixels of the image across several PM3b containing puncta and the ER network showed that the signal from PM3b puncta overlapped with the signal from the ER network tubules, junctions and cisternae (Fig. 1b).

**Fig. 1:**
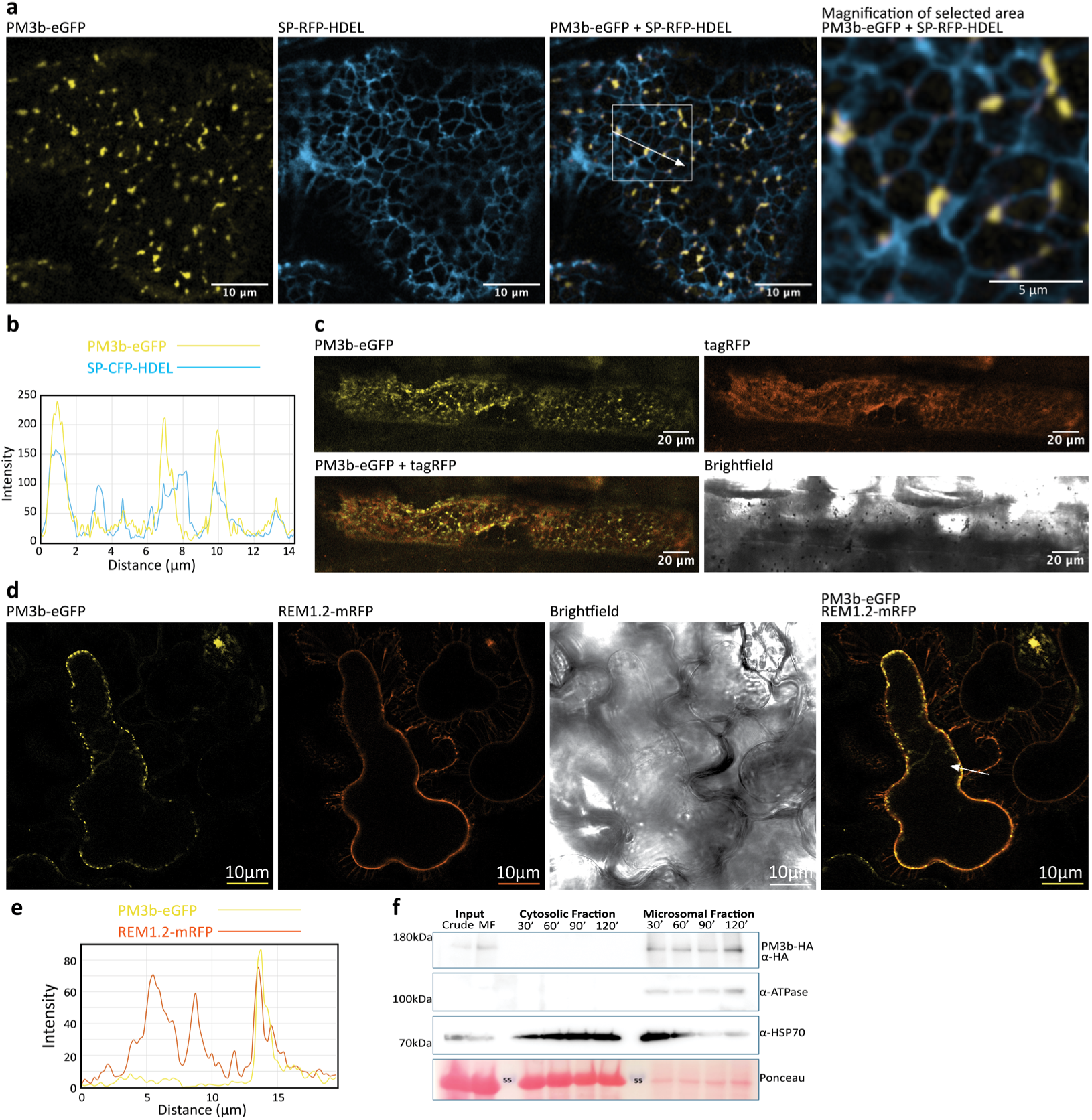
PM3b localizes to puncta at the endoplasmic reticulum network in *N. benthamiana* and wheat. (a) PM3b-eGFP co-localization with markers for the ER (SP-RFP-HDEL) in N. benthamiana. A magnified view of puncta associated with ER is shown as indicated by the selected area. (b) Pixel intensity measurements representing the fluorescence level taken across the white line depicted in (a). (c) PM3b-eGFP co-expression with the empty vector (EV) tagRFP which marks the cytoplasmic space in wheat. (d) PM3b-eGFP co-expressed with REM1.2-mRFP as a plasma membrane marker in N. benthamiana. Cells were incubated in a 4% NaCl hypotonic solution for 10-20 min before imaging. REM1.2-mRFP is present in the retracted cell membrane and the plasma membrane-cell wall contact sites of the Hechtian strands. PM3b-eGFP puncta retract from the cell wall as indicated by their absence in the Hechtian strands. (e) Pixel intensity measurements representing the fluorescence level taken across the white line depicted in (d). (f) Microsomal fractionation experiment of PM3b-HA showing its specific accumulation in the microsomal membrane fraction. Crude = total lysate and MF = membrane fraction. Images in (a) and (d) were taken 48-72 hours post infiltration using laser scanning confocal microscopy and 24-48 hours after bombardments in (c). Each confocal image represents a single plane of a cell. Scale bars are as indicated.

Localization patterns of a protein can vary depending on the species used for expression. *N. benthamiana* and wheat diverged from each other 200 million years ago (Humphry et al., 2010) and PM3b from wheat might therefore not localize in the same way in *N. benthamiana*. To investigate the localization pattern of PM3b in wheat, we transiently expressed PM3b-eGFP under a ubiquitin promoter in wheat epidermal cells through particle bombardments. For easier detection of transformed cells, we co-bombarded PM3b-eGFP with a vector that expresses tagRFP alone as a nucleocytoplasmic marker. LSCM revealed that PM3b-eGFP also localized to the puncta in wheat (Fig. 1c) as we had observed in *N. benthamiana*. Since PM3b showed a comparable localization pattern in wheat and *N. benthamiana* but we observed a low transformation rate in wheat bombardments, we performed further analysis of the puncta in the heterologous *N. benthamiana* system.

Next, we tested if PM3b-eGFP showed any colocalization with the plasma membrane (PM) marker REM1.2-mRFP (Bücherl et al., 2017). Transient expression of the two proteins together showed a bead on a string structure for PM3b which partially overlapped at the region marking the PM as indicated by the signal from REM1.2-mRFP (Supplementary Fig. 2a). To determine if PM3b is physically connected with the PM in *N. benthamiana*, we performed plasmolysis experiments in the presence of PM3b-eGFP and REM1.2-mRFP. During plasmolysis, REM1.2-mRFP was retracted from the cell wall and outlined the shrunken cytoplasm as well as long filaments of plasma membrane known as Hechtian strands which are still physically connected to the cell wall (Lang-Pauluzzi, 2000) (Fig. 1d). In contrast PM3b-eGFP remained in the puncta which were contained within the shrunken cytoplasm of the plasmolyzed cell and could not be found in the Hechtian strands (Fig. 1d & e), indicating that it is not integrated into or strongly associated with the PM but localized peripherally to it. Detailed analysis of the localization of the ER marker SP-CFP-HDEL, the PM marker REM1.2-mRFP and PM3b-eGFP during co-expression showed that PM3b was found near both the ER and PM (Supplementary Fig. 2a), suggesting that PM3b is not luminal to the ER but could be positioned at the outer surface of the ER network.

Due to its partial co-localization with ER and PM markers, we also tested if PM3b is associated with lipid membranes or exists freely in the cytoplasm in *N. benthamiana.* To do so, we performed sub-cellular fractionation experiments. We tested four different centrifugation times to pellet the microsomal fractions and found that already after 30 minutes, the PM was fully pelleted as indicated by the specific detection of the PM localized ATPase. We also detected HSP70 which can localize to the cytoplasm and ER as shown by its signal in both the supernatant and microsomal fractions at each centrifugation time. By expressing an HA tagged PM3b we found that it partitioned with the microsomal fraction which suggests that in *N. benthamiana*, PM3b physically associates with lipid microsomes (Fig. 1f). We conclude that PM3b localizes to puncta which are likely associated with the ER network in *N. benthamiana* and wheat.

### PM3b puncta are immobile in the plant cell and overlap with S-type ER-PM contact sites

The organelles within the cell display varying degrees of mobility over time. Mitochondria, vesicles derived from the Golgi network, and the ER network have high rates of diffusion while other structures remain relatively immobile (Christensen & Reck-Peterson, 2022). To better understand the sub-cellular localization of PM3b we tested whether PM3b puncta are immobile by transiently co-expressing PM3b-eGFP with the Golgi vesicle marker consisting of the first 49 amino acids of α-1,2-mannosidase I from soybean fused to mCherry (GmMAN1.2-mCherry) (Nelson et al., 2007) in *N. benthamiana*. LSCM revealed that PM3b-eGFP did not overlap with GmMAN1.2-mCherry (Fig. 2a) suggesting that the puncta are not contained within Golgi vesicles. To test for movement of Golgi vesicles and PM3b-eGFP, we performed LSCM time-lapse imaging. At timepoint 0 both the GmMAN1.2-mCherry marker and PM3b-eGFP were visible as distinct structures (Fig. 2b & c). An average overlay of the images in the time-lapse revealed that the Golgi-vesicle signal largely disappeared as it remained mobile, and spots would have changed position between frames. Conversely, an average overlay of the PM3b-eGFP puncta showed that they remained visible and were therefore more stationary than the Golgi vesicles (Fig. 2b & c). A max projection overlay of the time-lapse images of PM3b-eGFP in the same cell showed that the puncta were immobilized as they remained visible in the same distinct puncta over the entire time lapse. Again, the opposite was seen for the Golgi marker which due to its fast rate of diffusion created a smear of signal across the intracellular space (Fig. 2b & c). We conclude that the puncta of PM3b-eGFP which are present in the ER three-way junctions and tubules are immobilized, different to the highly dynamic Golgi vesicles.

**Fig. 2:**
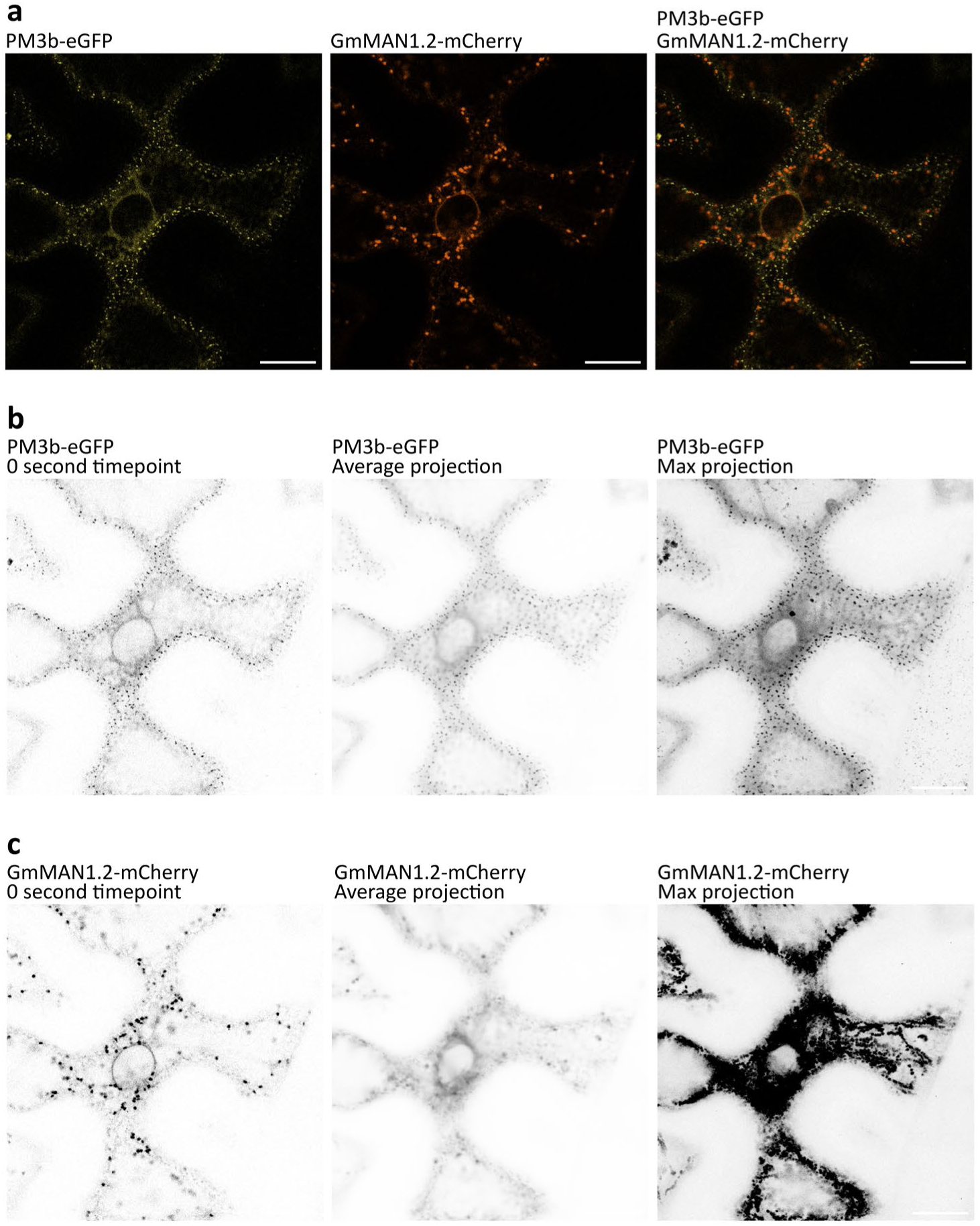
PM3b puncta are immobile in *N. benthamiana.* (a) PM3b-eGFP and the Golgi marker GmMAN1.2-mCherry marker were co-expressed in N. benthamiana. A merge between PM3b-eGFP and GmMan1.2-mCherry is also shown. (b & c) Inverted gray scale pictures of PM3b-eGFP and GmMAN1.2-mCherry to display movement of fluorescence signals. Left panel: timepoint 0s of PM3b-eGFP and GmMAN1.2-mCherry localization. Middle panel: average of signal intensity across entire timelapse imaging of PM3b-eGFP and GmMAN1.2-mCherry. Right panel: maximum of signal intensity (max projection or the sum of all signals) across the entire timelapse imaging of PM3b-eGFP and GmMAN1.2-mCherry . Images were taken 48-72 hours post infiltration using confocal laser scanning microscopy. Timelapse consisted of 139 individual images taken at 3-5 second intervals. Scale bar for all pictures is 20µm.

To maintain critical functional interactions with the PM, the ER is directly tethered to the PM at distinct contact sites. These ER-PM contact sites (EPCSs) are formed by specialized protein complexes that bring the regions of the ER and the PM within 10-30 nm of each other to allow for exchange of signaling molecules in a vesicle independent manner. Furthermore, these EPCSs are generally immobile and exist as puncta at the three-way junctions and cisternae of the ER network and the PM (Li et al., 2021; Perez-Sancho et al., 2015). In *Arabidopsis thaliana*, two well studied EPCS-localizing proteins are Synaptotagmin 1 (AtSYT1) and VAMP/Synaptobrevin-associated protein 27-1 (AtVAP27-1). It was shown that in both *A. thaliana* and *N. benthamiana*, they localize to distinct EPCS regions where AtVAMP27-1-enriched EPCSs (V-EPCSs) are always in contact with AtSYT1-enriched EPCSs (S-EPCSs). In the same study, they found that AtSYT1 labelled S-EPCS can be found flanking the enlarged V-EPCSs but also exist without contacting them in puncta along the ER network (Siao et al., 2016).

As described above, our results show that PM3b localizes into immobile puncta at distinct regions of the ER network that are proximal to but not integrated into the PM. Based on these findings, we hypothesized that PM3b could be found within EPCSs and tested for this by co-localizing PM3b with markers for S- and V-type EPCSs. To do this, we cloned AtSYT1 and AtVAP27-1 from *Arabidopsis thaliana* and tagged them with C-terminal tagRFPs. We co-expressed PM3b-eGFP and the ER marker SP-CFP-HDEL with either AtSYT1-tagRFP or AtVAP27-1-tagRFP in *N. benthamiana* and examined cells that were transformed with all three constructs via LSCM. We observed that AtVAP27-1 labelled V-EPCSs distributed into large puncta that were also in contact with the ER network (Fig. 3a). Furthermore, we found that the V-EPCS puncta showed an increased size with the overexpression of AtVAP27-1 as previously described (Siao et al., 2016). Upon co-expression with AtVAP27-1, we found that PM3b-eGFP puncta flanked the enlarged VAP27-1 labelled V-EPCSs while also existing in V-EPCS-free sites along the ER network (Fig. 3a & b). In line with these findings, co-expression of PM3b-eGFP and AtSYT1-tagRFP showed that they co-localized, as further substantiated by the intensity measurements across several of these puncta (Fig. 3c & d).

**Fig. 3:**
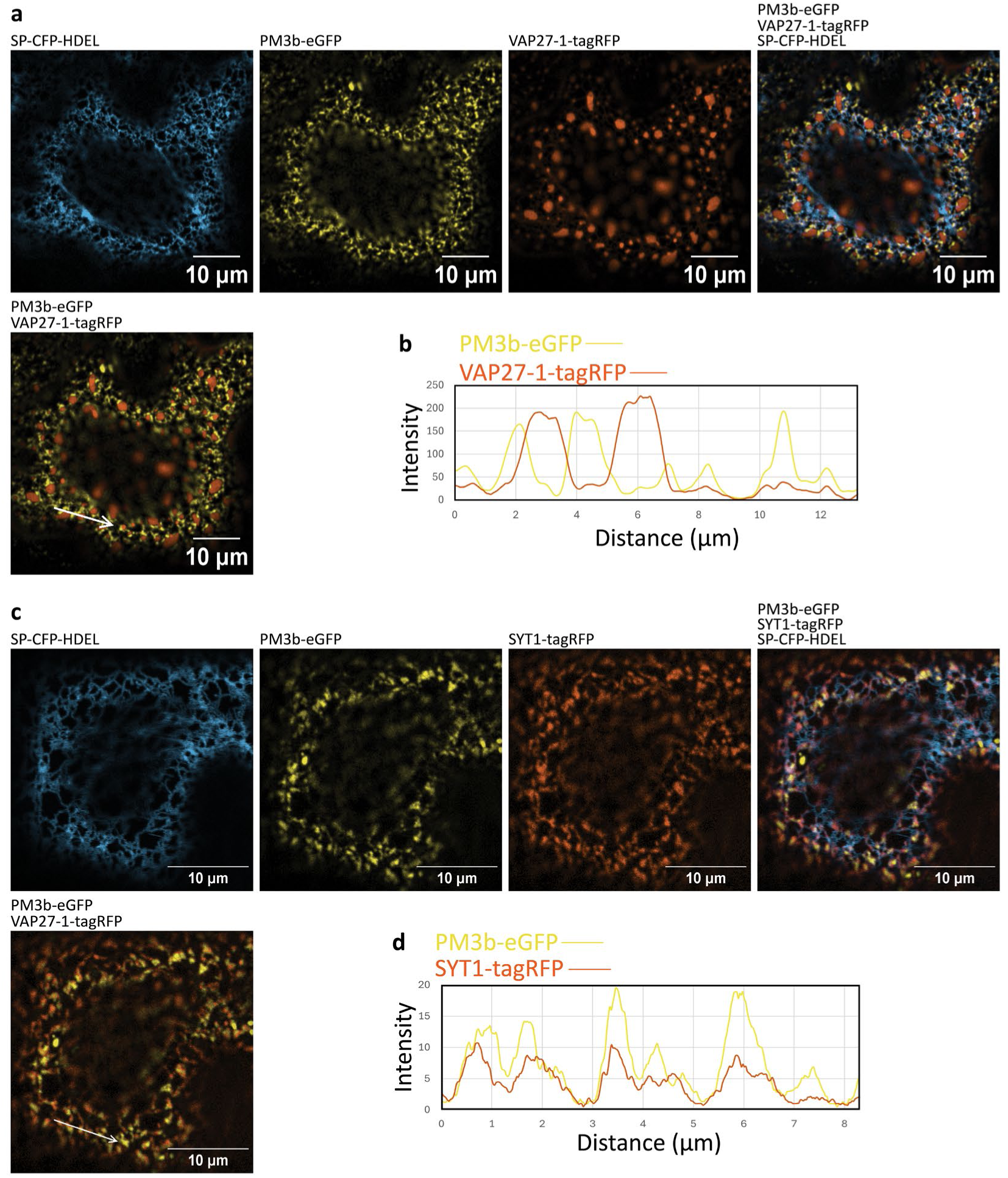
PM3b localizes to S-type ER-PM contact sites in *N. benthamiana*. (a) Images of PM3b-eGFP co-expressed with markers for the ER (SP-CFP-HDEL) and the V-EPCS marker VAP27-1-tagRFP are shown individually. Images of merges of PM3b-eGFP with both markers SP-CFP-HDEL and VAP27-1-tagRFP or PM3b- eGFP with the marker VAP27-1-tagRFP are also shown. Pictures were taken 48-72 hours post infiltration. (b) intensity measurements of the signal along the line of PM3b-eGFP and VAP27-1-tagRFP, indicating that the signals do not overlap significantly but flank each other. (c) Images of PM3b-eGFP co-expressed with markers for the ER (SP-CFP-HDEL) and the S-EPCS marker SYT1-tagRFP are shown individually. Images of merges of PM3b-eGFP both markers SP-CFP-HDEL and SYT1-tagRFP or PM3b-eGFP with the marker SYT1-tagRFP are also shown. Pictures were taken 48-72 hours post infiltration. (d) intensity measurements of the signal along the line of PM3b-eGFP and SYT1-tagRFP indicate that PM3b-eGFP and SYT1-tagRFP signals overlap with each other. Images in A and C were taken 48-72 hours post infiltration using confocal laser scanning microscopy. Each confocal image represents a single plane of a cell. Scale bars are as indicated.

These results suggest that in its resting state, PM3b-eGFP puncta are co-localizing with AtSYT1 into immobile structures at the ER network that represent the specific subset of EPCSs known as S-EPCSs.

### AVRPM3^b2/c2^ localizes to the cytoplasm and nucleus in different ratios in a variant dependent manner

The two active AVR variants that are recognized by PM3b are AVRPMB3^b2/c2^-A and AVRPM3^b2/c2^-I. Variant I of AVRPM3^b2/c2^ has a single amino acid polymorphism from a serine (S) to an asparagine (N) in position 21 of the mature peptide as compared to the A variant. In a comparison of the strength of HR in *N. benthamiana*, AVRPM3^b2/c2^-A elicited HR with PM3b and weak HR with PM3c while AVRPM3^b2/c2^-I elicited HR with both PM3b and PM3c and the HR accumulated faster than for AVRPM3^b2/c2^-A, which designated it as the stronger variant (Bourras et al., 2019). To test the localization of these active variants, we fused both variant A and I of AVRPM3^b2/c2^ C-terminally with eGFP (AVRPM3^b2/c2^-A-eGFP and AVRPM3^b2/c2^-I-eGFP) and expressed them in *N. benthamiana*. As a large fusion protein such as eGFP might affect the function of a protein, we tested the ability of AVRPM3^b2/c2^-A-eGFP to elicit HR in the presence of PM3b-MYC in *N. benthamiana* and we detected no HR. When we tested AVRPM3^b2/c2^-I-eGFP with PM3b-MYC we found that there was a detectable amount of HR although this was lower than with AVRPM3^b2/c2^-I-HA when combined with PM3b-MYC (Supplementary Fig. 1b).

LSCM in *N. benthamiana* showed that both AVRPM3^b2/c2^-A-eGFP and AVRPM3^b2/c2^-I-eGFP localized to the nucleocytoplasmic space (Fig. 4a & b) However, the signal of AVRPM3^b2/c2^-I-eGFP in the nucleus was higher than that of AVRPM3^b2/c2^-A-eGFP as measured by the fluorescence intensity across the nucleus and the cytoplasmic space directly surrounding it (Fig. 4c & d). The elevated nuclear localization of AVRPM3^b2/c2^-I-eGFP could have been due to a higher amount of cleavage product. However, immunoblotting of protein lysates from *N. benthamiana* transiently expressing AVRPM3^b2/c2^-A-eGFP or AVRPM3^b2/c2^-I-eGFP showed that no significant amount of cleaved eGFP was present (Fig. 4e & f). To conclude, the two recognized AVRPM3^b2/c2^ variants show similar sub-cellular localization patterns to the nucleocytoplasmic space but AVRPM3^b2/c2^-I shows a stronger localization to the nucleus.

**Fig. 4:**
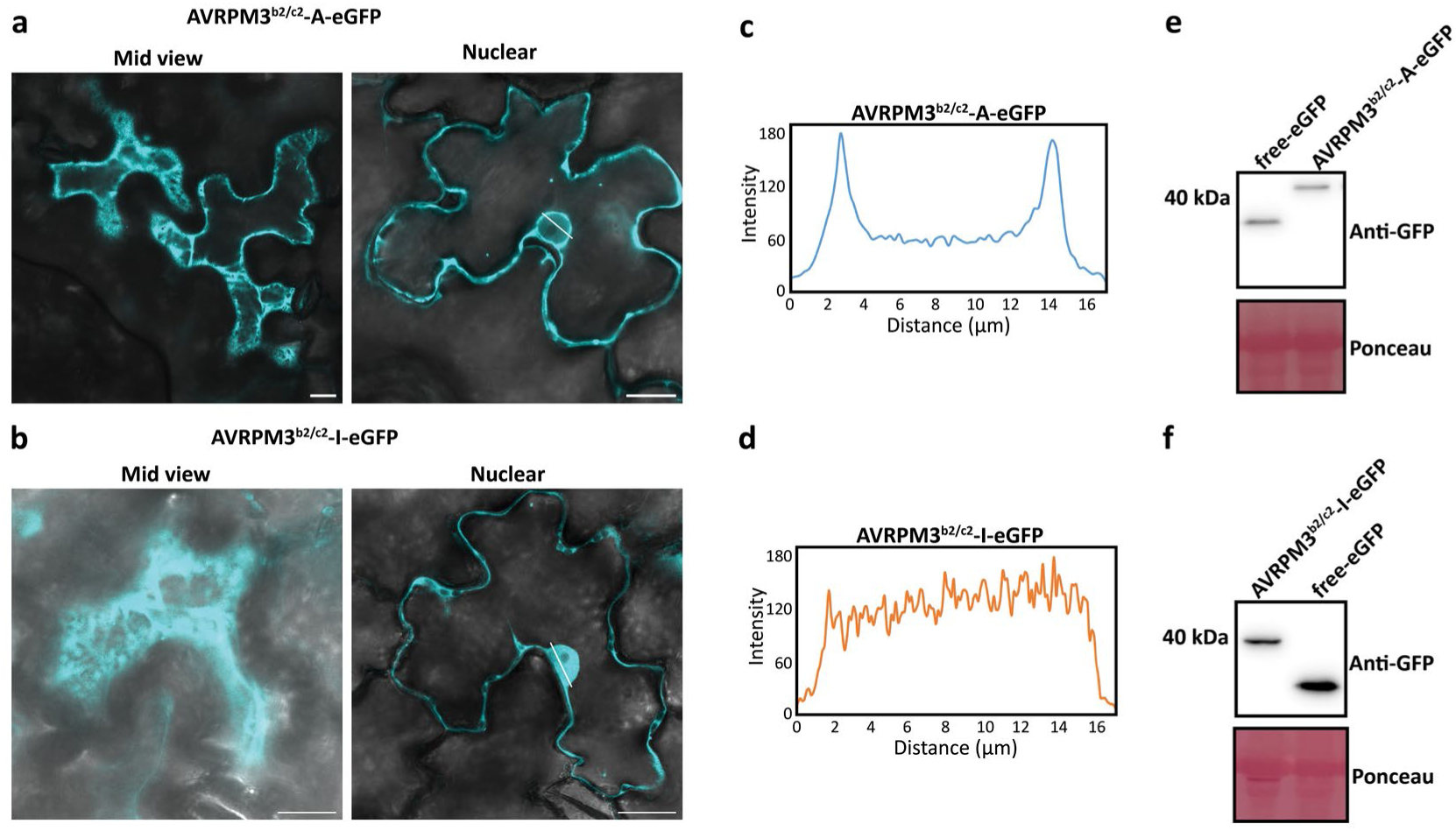
Localization of different AVRPM3^b2/c2^ variants. (a) View of the cytoplasmic strands (mid view) and the nucleus (nuclear) in the same N. benthamiana cell showing differential nucleocytoplasmic localization patterns of (a) AVRPM3^b2/c2^-A-eGFP or (b) AVRPM3^b2/c2^-I-eGFP. Pictures were taken using confocal laser scanning microscopy 48-72 hours post infiltration. (c & d) Measurement of the fluorescence intensity across the nucleus for AVRPM3^b2/c2^-A-eGFP and AVRPM3^b2/c2^-I-eGFP. The measurement was made for both AVR variants across the white line drawn across the nucleus in pictures (a) and (d), respectively. (e and f) Immunoblots probed with anti-GFP-HRP antibodies show protein accumulation of variants AVRPM3^b2/c2^-A-eGFP and AVRPM3^b2/c2^-I-eGFP in the localization studies and free eGFP as a control. Loading control was done with Ponceau staining of each immunoblot, displaying equal levels of the rubisco large subunit. Scale bar for all images is 20µm.

### AVRPM3^b2/c2^ interacts with the LRR domain of PM3b in planta

The binding mechanism between PM3 allelic variants and their corresponding AVRPM3 effectors is not known. The expression of PM3b with AVRPM3^b2/c2^ in *N. benthamiana* leads to a HR and cell death, suggesting a well conserved mechanism across plant species (Bourras et al., 2019). The localization of PM3b in distinct puncta in the cell, corresponding to EPCSs, suggests that this could also be the site of interaction with AVRPM3^b2/c2^. However, AVRPM3^b2/c2^ is only observed in the cytoplasm and nucleus. For a direct interaction between the proteins or a complex formation in an indirect interaction, PM3b could still be localizing to the outer cytoplasmic face of the ER network which would allow for interaction with AVRPM3^b2/c2^. Therefore, we also wanted to test if PM3b interacts with AVRPM3^b2/c2^ in planta.

First, we tested the effect of co-expression of PM3b with variant A of AVRPM3^b2/c2^ to establish a timepoint at which we could detect both proteins before a strong cell-death response and thus optimize protein expression levels. To do so, we co-expressed PM3b-HA with AVRPM3^b2/c2^-A-FLAG or either one alone in *N. benthamiana* and collected samples at 24 hours post infiltration (hpi), 48 hpi and 72 hpi. To have comparable expression levels of PM3b-HA and AVRPM3^b2/c2^-A-FLAG when expressed separately, we substituted their co-expressor with GUS (Bourras et al., 2015). Immunoblotting of AVRPM3^b2/c2^-A-FLAG co-expressed with PM3b-HA revealed that low but detectable amounts of AVRPM3^b2/c2^-A-FLAG were found at 48 hpi at a slightly lower level than for AVRPM3^b2/c2^-A-FLAG with GUS (Fig. 5a). However, at 72 hpi, AVRPM3^b2/c2^-A-FLAG was no longer detectable when co-expressed with PM3b-HA, while protein levels of AVRPM3^b2/c2^-A-FLAG with GUS kept increasing (Fig. 5a). In the two samples PM3b-HA with AVRPM3^b2/c2^-A-FLAG and PM3b-HA with GUS, immunoblotting of PM3b-HA showed low but similar levels of protein accumulation already at 24 hpi (Fig. 5a). Furthermore, PM3b-HA protein continued to accumulate at similar levels in both samples at 48 hpi and 72 hpi. This suggests that PM3b-HA co-expression with AVRPM3^b2/c2^-A-FLAG did not modify the levels of PM3b-HA over the 72-hour time-course but the reduced protein level was specific to AVRPM3^b2/c2^-A when PM3b was present (Fig. 5a). When PM3b is co-expressed with AVRPM3^b2/c2^-A, cell death only visually occurs 4-5 days post infiltration and our samples did not show any visual cell death phenotype during our time course experiment. We conclude that for co-expression and subsequent co-immunoprecipitation experiments, collection at 24-36 hpi will lead to the highest accumulation of both proteins and that the reduction in AVRPM3^b2/c2^ protein levels is not due to a strong cell death response but is likely intrinsic to the presence of and possibly the interaction with and activation of PM3b.

**Fig. 5:**
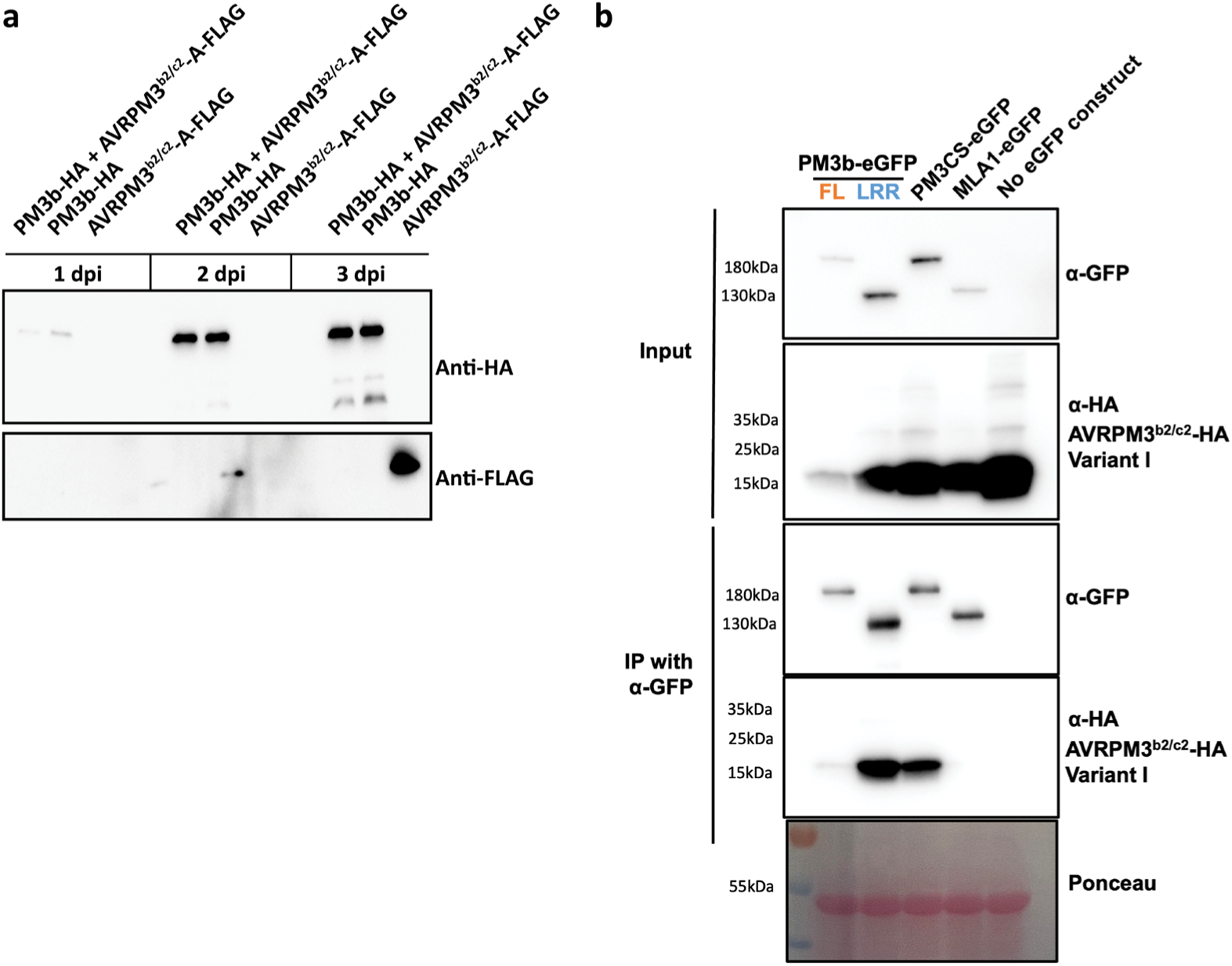
AVRPM3^b2/c2^ interacts in planta with PM3b and PM3CS through the LRR domain. (a) Time course experiment to test for the expression of AVRPM3^b2/c2^-A and PM3b alone or co-expressed. AVRPM3^b2/c2^ shows reduced accumulation in the presence of PM3b. Immunoblots were probed with α-HA, α-FLAG or α-GFP antibodies as indicated. (b) Co-immunoprecipitation experiment using α-GFP agarose beads of HA tagged avirulent variant I of AVRPM3^b2/c2^ with eGFP tagged full length (FL) PM3b, its LRR domain, PM3CS, MLA1 or without an eGFP tagged construct.

Next, we performed co-immunoprecipitation to establish potential interaction of PM3b with the AVRPM3^b2/c2^ effector protein. To have the strongest possible interaction response with PM3b in co-immunoprecipitation experiments, we used the strongly recognized variant I instead of variant A of AVRPM3^b2/c2^, which was previously shown to induce the strongest PM3b-mediated HR response among all naturally occurring AVRPM3^b2/c2^ variants (Bourras et al., 2019). We co-expressed an HA-tagged version of the strong HR inducing AVRPM3^b2/c2^-I with eGFP tagged full length (FL) PM3b or its LRR domain (PM3b-LRR) alone in *N. benthamiana* followed by co-immunoprecipitation. To test for specificity, we used the eGFP tagged susceptible variant PM3CS (Yahiaoui et al., 2006) and the barley CC-NLR MLA1 which gives resistance to the barley powdery mildew pathogen *Blumeria hordei* (*Bh*) (Lu et al., 2016) by directly recognizing AVRA1 (Saur et al., 2019). We found that the amount of AVRPM3^b2/c2^-I present in the sample with the FL PM3b was lower than with PM3b-LRR or with PM3CS (Fig. 5b). Furthermore, the level of FL PM3b was slightly lower as compared to PM3CS levels but we enriched and recovered similar amounts of both proteins in our immunoprecipitations (Fig. 5b). Interestingly, we found that AVRPM3^b2/c2^-I interacted only weakly with FL PM3b but showed a stronger interaction with PM3b-LRR and PM3CS (Fig. 5b). The interaction of AVRPM3^b2/c2^-I with FL PM3b, PM3b-LRR and PM3CS was specific as AVRPM3^b2/c2^-I did not interact with MLA1 or the α-GFP agarose beads alone (Fig. 5b).

We conclude that PM3b can interact with AVRPM3^b2/c2^ and the LRR domain alone is sufficient for this interaction. Furthermore, our observations suggest that AVRPM3^b2/c2^ associates with PM3b through amino acids shared between different PM3 alleles such as PM3CS, and the PM3b specificity for full activation is determined within the residues distinct to PM3b.

### Activation of PM3b releases AVRPM3^b2/c2^ association

PM3b and PM3CS are the two most divergent PM3 variants yet still show a remarkable 97% identity. Furthermore, their CC domains are identical and the vast majority of the amino acid polymorphisms between PM3b and PM3CS are found in the NB-ARC domain. In the LRR domain, PM3b differs from PM3CS by only three amino acids: L587I, the insertion R588 in the N-terminal region and I1309M in the C-terminal region (Yahiaoui et al. 2006). AVRPM3^b2/c2^interacts with PM3b-LRR and PM3CS but when co-expressed with PM3b, the protein level of AVRPM3^b2/c2^ is reduced and the interaction is weaker (Fig. 5b). We hypothesize that the activation of PM3b leads to a disassociation from AVRPM3^b2/c2^. Therefore, AVRPM3^b2/c2^ should interact to a similar extent with an inactive PM3b as it does with PM3CS. NLR activation has been shown to be abolished when mutating the ATP binding site located in the P-loop, also known as the Walker A motif. This ATP binding site has the consensus amino acid sequence GGLGKTTL (He et al., 2022) which is also preserved in PM3b in position 208-215. To create an inactive version of PM3b, we generated the K212A (PM3b^K212A^) P-loop mutant. We tested if eGFP tagged PM3b^K212A^ was non-functional, and as expected it did not induce cell death when co-expressed with AVRPM3^b2/c2^-I (Supplementary Fig. 1c). In addition to using the inactive PM3b^K212A^ we also tested if the virulent variant C of AVRPM3^b2/c2^ (AVRPM3^b2/c2^-C) would show a stronger interaction with PM3b as compared to the avirulent AVRPM3^b2/c2^-I (Bourras et al., 2019). To do so, we co-expressed HA-tagged versions of variant C and I of AVRPM3^b2/c2^ with eGFP tagged versions of the NLRs in *N. benthamiana* and performed co-immunoprecipitation experiments. This revealed a robust interaction of PM3b^K212A^ with both variants of AVRPM3^b2/c2^ which was as strong as PM3CS interaction with both AVRPM3^b2/c2^ variants (Fig. 6a). In contrast, only the virulent AVRPM3^b2/c2^-C interacted with PM3b to a level similar to that of PM3CS and PM3b^K212A^ interaction with both AVRPM3^b2/c2^ variants. To further validate these results, we performed the reciprocal co-immunoprecipitation experiment using HA-tagged AVRPM3^b2/c2^ as the bait protein and eGFP tagged NLRs as the prey. Here again we observed a strong interaction between PM3b^K212A^ and both the I- and the C-variants of AVRPM3^b2/c2^ while PM3b only interacted with AVRPM3^b2/c2^-C to a similar extent as PM3b^K212A^ (Fig. 6b). We wondered whether the different PM3 variants would have the same sub-cellular localization, as this could be affecting their function. LSCM of cells expressing PM3b^K212A^-eGFP and PM3CS-eGFP in *N. benthamiana* showed that both proteins localized similarly to PM3b to the S-EPCS localized puncta (Supplementary Fig. 3a). This strongly suggests that a shared feature in PM3 allelic variants is responsible for their localization.

**Fig. 6:**
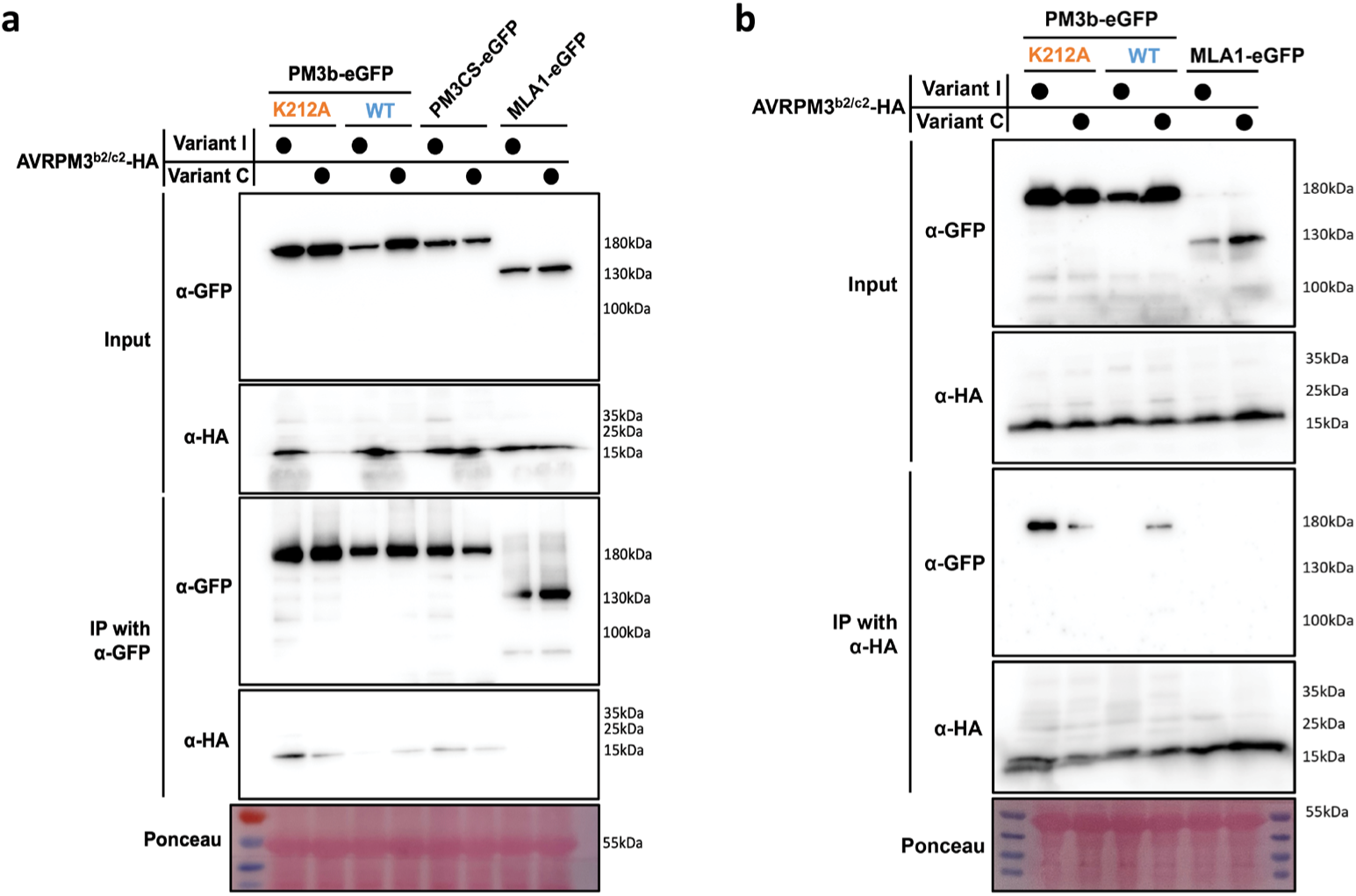
PM3b activation leads to a transient interaction with AVRPM3^b2/c2^. (a) Immunoprecipitation of eGFP tagged P-loop mutant PM3b^K212A^, wildtype PM3b, PM3CS or MLA1 transiently co-expressed with HA tagged avirulent variant I or virulent variant C of AVRPM3^b2/c2^. (b) Immunoprecipitation of HA tagged avirulent variant I or virulent variant C of AVRPM3^b2/c2^ transiently co-expressed with eGFP-tagged P-loop mutant PM3b^K212A^, wildtype PM3b or MLA1. Each blot was stained with Ponceau to show equal loading between samples. Immunoblots were probed with α-HA or α-GFP as indicated.

We conclude that the strength of the interaction between PM3b and AVRPM3^b2/c2^ variants is inverse to the strength of the HR in *N. benthamiana*. Here, the stronger interaction with PM3b is seen for the virulent AVRPM3^b2/c2^-C, which elicits no HR, than for avirulent and strong HR inducing AVRPM3^b2/c2^-I. This suggests a transient association between the two proteins where upon activation of PM3b, its ability to bind AVRPM3^b2/c2^ is abolished.

## Discussion

In this study, we describe a novel sub-cellular localization of an NLR immune receptor: in its resting state, PM3b localizes at an important signaling interface between the ER and the PM at immobile membrane contact sites. In contrast, its corresponding avirulence effector AVRPM3^b2/c2^ localizes mainly to the cytoplasm and nucleus. Nonetheless, we could detect an interaction between PM3b and AVRPM3^b2/c2^ after co-infiltration in *N. benthamiana*, suggesting that PM3b localization is to the outer surface of the ER membrane which would allow it to interact with AVRPM3^b2/c2^. We found that active PM3b did not strongly interact with AVRPM3^b2/c2^ while inactivation of PM3b through a mutation in its P-loop stabilized this interaction. Furthermore, we could show that the susceptible allelic variant PM3CS interacted with AVRPM3^b2/c2^ to a similar extent as the P-loop mutant of PM3b. Together this would suggest that AVRPM3^b2/c2^ interacts with PM3 variants through amino acids which are conserved between PM3b and PM3CS while full activation of PM3b is determined by amino acids specific to this variant.

Diverse sub-cellular localization including the cytoplasm, nucleus, plasma membrane, tonoplast and endoplasmic reticulum have been described in localization studies of NLR-type immune receptors (Lolle et al., 2020; Lüdke et al., 2022; Qi & Innes, 2013; Sheperd et al., 2023). Often, the localizations of immune receptors are matched to the sites of their corresponding AVR’s localization and function, and in some cases, they are re-localized upon AVR presence (Deslandes et al., 2003; Du et al., 2015; Duggan et al., 2021). We showed that PM3b-eGFP resides in immobile puncta that co-localize with the ER network. Interestingly, other NLR proteins in wheat and the sister species barley show different localizations as compared to PM3b. MLA proteins and their wheat ortholog, the stem rust immune receptor SR50, localize mainly to the cytoplasm and nucleus in barley and wheat, respectively, as well as in *N. benthamiana* (Ortiz et al., 2022; Shen et al., 2007). In another case, the stem rust immune receptor, SR35 was shown to localize to the PM (Zhao et al., 2022). The wheat powdery mildew immune receptor, PM2 was suggested to indirectly recognize AVRPM2 through a complex interaction with a putative effector target, the wheat zinc finger protein TaZF. While PM2 was mainly found in the cytosol, interaction with TaZF recruited it and AVRPM2 to the nucleus (Manser et al., 2024). Contrastingly, PM3b associates with lipid microsomes at ER network puncta which seems to be a common feature of PM3 immune receptors.

The structure of the ER is divided into several sheets called cisternae, tubules, and other smaller domains that work together to allow it to function correctly (English et al., 2009). In eukaryotes, a unique feature called membrane contact site have been discovered which have been shown to allow close inter-organelle contacts, dissipation of various signals and exchange of material (Prinz et al., 2020). The PM for example can form a close association with the ER through ER-PM contact sites (EPCS) which are facilitated by various tethering proteins (Pérez-Sancho et al., 2015). The powdery mildew-plant interface contains an extra-haustorial membrane (EHM) derived from the plant that encases the developing haustoria and shares properties with the ER (Kwaaitaal et al., 2017). Furthermore, the EHM has also been shown to have EPCS like connections that can potentially bring proteins such as SYT1 and PEN1, which have antagonistic effects on *A. thaliana* immunity to mildew, into close contact with the interface (Ishikawa et al., 2018; Siao et al., 2016). The co-localization of the PM3b puncta with the ER junctions and tubules suggests that they are closely associated with it. EPCS exist in at least two different forms of either VAP27-1 labelled (V-EPCS) or SYT1 labelled (S-EPCS) (Siao et al., 2016). Our study shows that PM3b co-localizes with S-EPCSs while it was excluded from V-EPCSs. This showed that PM3b localization is highly specific and overlaps with components involved in immunity against powdery mildew such as SYT1.

In a recent study, Contreras and colleagues (2023) investigated the activation of the helper NLR required for cell death (NRC) through its sensor NLR Rx which confers resistance to *Potato virus X* (PVX) in potato (*Solanum tuberosum*) by recognizing the coat protein (CP) as its AVR. In this work, it was found that upon activation of Rx, downstream signaling activates NRC2, which oligomerizes into a resistosome that does not include Rx. Notably, the NRC2 oligomer associates with the plasma membrane in puncta reminiscent of the puncta observed for PM3b. In another study, it was found that the helper-NLR NRC4 dynamically relocates from the cytoplasm to the EHM of *Phytophthora infestans* during infection while NRC2 did not. Upon activation in the presence of its sensor-NLR Rpi-blb2, NRC4 also forms puncta which spread from the focal accumulation around the EHM to also include the PM (Duggan et al., 2021). As PM3b accumulates in puncta already in a resting state, this differs from what has been observed for NRC2 and NRC4 which only form puncta upon activation (Contreras et al., 2023; Duggan et al., 2021). Until now, our attempts at localizing PM3b in an active state has failed as the signal was abolished when using the autoactive PM3b^D502V^-eGFP or when co-expressing PM3b-eGFP with AVRPM3^b2/c2^, possibly due to the death of most of the expressing cells or events leading to the cell death response. Thus, it remains open if the localization to the puncta is important for cell death signaling. One possible reason for PM3b localization to S-EPCS could be to allow for early detection of AVRPM3^b2/c2^ during *Bgt* infection. SYT1 has been shown in oomycetes to accumulate around the haustoria in the EHM (Duggan et al., 2021). Since PM3b and SYT1 co-localize into S-EPCS, both could potentially be recruited to the EHM of *Bgt* which would allow for early detection of AVRPM3^b2/c2^ when it is secreted into the plant cell.

Our initial co-immunoprecipitation experiments between PM3b and AVRPM3^b2/c2^-I revealed that the two proteins interacted, and this interaction is likely to be mediated by the LRR domain of PM3b (Fig. 5b). The LRR domain of PM3 is made up of 29 repeats and a comparison of functional variants of PM3 showed that the predicted solvent exposed residues of the LRR are undergoing diversifying selection which suggests that the LRR is involved in recognition of its corresponding AVRPM3 effector (Srichumpa et al., 2005). In a recent study, the structure of MLA13-AVRa13-1 was solved as a heterodimer complex by cryo-EM (Lawson et al., 2024). The study showed that AVRa13-1 is an RNase-like effector which binds through multiple contact points to the C-terminal LRRs and winged helix domain (WHD) of MLA13. A previous study on PM3 variant specificities showed that an introduction of the amino acid change I1309M into PM3CS from PM3b is seemingly enough to lead to a partial activation (Brunner et al., 2010). Strikingly, we could also detect an interaction between the susceptible variant PM3CS and AVRPM3^b2/c2^-I which was stronger than the one with PM3b. This would indicate that PM3b and PM3CS interact with AVRPM3^b2/c2^ through conserved residues between the two variants and the amino acids unique to PM3b determine its ability to active a cell death response. The weaker interaction we observed with PM3b as compared to the LRR domain of PM3b or with PM3CS suggested that activation of PM3b leads to dissociation from AVRPM3^b2/c2^. This was further substantiated when we found that a strong interaction of PM3b with AVRPM3^b2/c2^ could be re-constituted by the P-loop mutation K212A in PM3b^K212A^, rendering it unable to bind ATP and become fully activated (Fig. 6a). Similarly, when we used the virulent variant AVRPM3^b2/c2^-C which does not elicit HR, it could still interact with PM3b (Fig. 6b). In contrast, MLA, SR35 and SR50 show a direct interaction with their avirulence effector targets and stay stably bound in a complex upon NLR activation (Ortiz et al., 2022; Saur et al., 2019; Zhao et al., 2022). It is currently unknown whether PM3b forms a resistosome upon activation. However, based on our findings we hypothesize that if PM3b does form a resistosome, this is likely to be in a confirmation excluding AVRPM3^b2/c2^. It is also unknown at this point if activation of PM3b leads to dissociation from S-EPCSs or if this is the final site of downstream signaling and cell death. Furthermore, PM3CS localized to similar puncta as PM3b (Supplementary Fig. 3a) indicating that this is also a shared feature between PM3 variants. Future work will complement our findings by studying the other PM3 variants and their AVRPM3 effector targets as well as the localization of PM3b or other PM3 variants during *Bgt* infection in wheat. The recent boom in structural studies of NLR-AVR complexes along with the knowledge gained from this study about interaction dynamics of PM3b and AVRPM3^b2/c2^ will facilitate the design of future experiments to study structure-function relationships between PM3 and AVRPM3 effector pairs.

## Materials and Methods

### Cloning of Arabidopsis thaliana genes

*AtSyt1*_AT2G20990.3_CDS and *AtVap27-1*_AT3G60600.1_CDS from TAIR (https://www.arabidopsis.org/) were used to design primers excluding the stop codon of the gene and including attB-site overhangs for BP cloning using Gateway® BP Clonase™ II Enzyme Mix (Thermo Fisher Scientific, Waltham, MA, USA). Sequences for *AtSyt1* and *AtVap27-1* amplified and used for further cloning are listed in supplementary table 1. Sequences were amplified from *A. thaliana* Col-0 cDNA using a homemade Pfu7x polymerase. Amplicons with BP cloning overhangs were inserted into pDONR207 using a BP Clonase II enzyme (Thermo Fischer Scientific, Waltham, MA, USA).

### Site directed mutagenesis

To mutate the P-loop in PM3b, we mutated Lysine in position 212 to an Alanine. Whole plasmid amplification of pENTR containing the cDNA sequence of *Pm3b* was performed with non-overlapping PCR primers and homemade Pfu7x polymerase. Amplicon was purified with a plasmid purification kit (Qiagen) followed by phosphorylation of linearized PCR amplicon using T4 polynucleotide kinase (New England Biolabs) and ligation with T4 ligase (New England Biolabs) according to manufacturer’s recommendations.

### Plasmid cloning

Sequences for *Avr* and *Nlr* constructs used in this study can be found in supplementary table 1. Vectors used in this study were pGWB505, pGWB228, pGWB235 (Nakagawa et al., 2009), pIPKB004 (Himmelbach et al., 2007) and pAHC17 (Christensen & Quail, 1996). C-terminal HA and FLAG tagged *AvrPm3^b2/c2^* constructs and C-terminally HA and MYC tagged *Pm3b* construct were cloned into the 35S promoter driven binary vector pIPKB004 (Himmelbach et al., 2007) as previously described (Bourras et al., 2019). C-terminal fluorescent protein fusions were created by cloning the sequence of interest from a Gateway® entry vector into the Gateway® destination vectors pGWB505 (eGFP) or pGWB228 (tagRFP) (Nakagawa et al., 2007) using Gateway® LR Clonase II based cloning (Thermo Fischer Scientific, Waltham, MA, USA). For expression in wheat, *Pm3b-eGFP* was amplified out of pGWB505-Pm3b and inserted into pAHC17 using the KpnI and SpeI restriction sites and enzymes (New England Biolabs) and T5 exonuclease-dependent assembly (TEDA) (Xia et al., 2019).

### Transient expression in *Nicotiana benthamiana*

*Agrobacterium tumefaciens* transformed with binary vectors containing the construct of interest were prepared in Luria broth (LB) containing the appropriate antibiotics and incubated overnight at 28°C with 200rpm shaking. Overnight cultures were harvested by centrifugation at 3,300xg for 5 minutes and washed by re-suspending in fresh LB media without antibiotics. Cells were harvested again by centrifugation at 3,300xg for 5 minutes and re-suspended in a small amount of AS-medium (10mM MES-KOH, pH5.6; 10mM MgCl_2_; 150µM acetosyringone) and diluted to an OD of 1.0 followed by 2-4 hours of incubation at 28°C with 200rpm shaking to induce virulence. Infiltrations into *Nicotiana benthamiana* were done by mixing the prepared *A. tumefaciens* cultures in the indicated ratios in the methods sections below.

### Transient expression in wheat through particle bombardment

pAHC17-Pm3b-eGFP and pGWB235-empty were used for Particle bombardment of single wheat epithelial cells which was performed as described previously (Brunner et al., 2010) with minor adjustments. 7–9-day old primary leaves of the wheat cultivar Bobwhite were bombarded with 7μg of each construct and incubated for 24-48 hours at 20°C and 80% relative humidity prior to imaging.

### Protein extraction, time-course and Western blot analysis

Proteins were transiently expressed in *N. benthamiana* using Agrobacterium mediated transformation in a construct:p19 ratio of 1:1 where p19 is an Agrobacterium p19-silencing-suppressor strain (Voinnet et al., 2003) and 3 days post infiltration, 6 leaf discs (6mm diameter) were collected. For time-course analysis, proteins were expressed in a PM3b:AVRPM3^b2/c2^:p19 ratio of 1:4:1 or with either PM3b or AVRPM3^b2/c2^ replaced by GUS. 6 leaf discs (6mm diameter) per sample were collected at 1 day post infiltration (dpi), 2 dpi and 3 dpi. Samples were ground in liquid nitrogen after which 150µl of 2x Laemmli buffer (62mM tris-HCl pH6.8, 10% glycerol, 2% SDS, 0.02% bromophenol blue, 1% β-mercaptoethanol) was added and samples were incubated at 55°C for 10 minutes followed by centrifugation at 22,000xg for 5 min. The supernatant of the clarified whole cell lysate was separated on 4-20% SDS-PAGE and transferred to a PVDF membrane. Membranes were probed with HRP conjugated primary anti-GFP (sc-9996 HRP, Santa Cruz), HRP conjugated primary anti-HA (clone 3F10, 12013819001, Roche) or anti-FLAG (M2, F3165, Sigma-Aldrich) and secondary HRP conjugated anti-mouse (sc-516102, LabForce).

### Microsomal fractionation

Proteins were transiently expressed in *N. benthamiana* using Agrobacterium mediated transformation with a ratio of PM3b:p19 of 1:1. Microsomal fractionation was performed as previously described (Abas & Luschnig 2010) with 100mg of starting material for each construct combination. Samples were separated on 4-20% SDS-PAGE and transferred to a PVDF membrane. Membranes were probed with HRP conjugated primary anti-HA (clone 3F10, 12013819001, Roche), anti-ATPase (AS07 260, Agrisera) followed by HRP conjugated anti-rabbit (sc-2357, LabForce) or anti-HSP70.

### Co-immunoprecipitation

Proteins were transiently expressed in *N. benthamiana* using Agrobacterium mediated transformation in a ratio of NLR:AVR:p19 of 2:1:1. For MLA1:AVR:p19 and PM3b-LRR:AVR:p19 the ratio was 1:1:1. Tissue was collected 24-36 hours post infiltration and ground in liquid nitrogen followed by protein extraction from 350mg of ground tissue in 1.5ml of lysis buffer (50mM tris-Cl, pH8.0; 150mM NaCl; 5.0mM MgCl_2_; 2.0mM Na_2_S_2_O_5_; 10% glycerol; 5.0mM DTT; 1.0mM EDTA; 0.5% IGEPAL CA-630 (Sigma-Aldrich); 0.2% Triton X-100 (Sigma-Aldrich); 1mM PMSF; 1x Protease inhibitor cocktail EDTA free (Roche) for 30 minutes at 4°C and constant rotation. Lysate was clarified twice by centrifugation at 22,000xg for 15 minutes after which some of the supernatant was collected for input samples. 1.0ml of supernatant was mixed with 10µl of pre-washed PierceTM anti-HA (88837, Thermo Fisher scientific) magnetic beads or 14μl of anti-GFP (Chromotek) agarose beads and incubated for 2-3 hours at 4°C with constant rotation. Beads were washed 5 times for 5 minutes each with 1.0ml of lysis buffer for HA-magnetic beads or with lysis buffer supplemented with 150mM NaCl for a final concentration of 300mM NaCl for GFP-agarose beads. Elution was performed by mixing beads with 50µl of 2x Laemmli buffer and incubating at 55°C for 15 minutes at 1,000 rpm. Proteins from input and eluates were separated on 4-20% or 4-18% home-made SDS-PAGE and transferred onto PVDF membranes (GE Healthcare, Chicago, Illinois, USA) followed by detection using HRP conjugated primary anti-HA (clone 3F10, 12013819001, Roche) and HRP conjugated primary anti-GFP (sc-9996 HRP, Santa Cruz).

### Confocal laser scanning microscopy and plasmolysis

NLRs and AVRs were cloned into pGWB505 (C-terminal eGFP) as described in plasmid cloning section. Cytosolic marker was free tagRFP expressed by transforming *A. tumefaciens* with pGWB235-empty (Nakagawa et al., 2007). This allows for expression of “free” tagRFP with a short C-terminal amino acid tail (ITSLYKKAERET-STOP). The ER marker was SP-mRFP-HDEL (Nelson et al., 2007), the PM marker was REM1.2-mRFP (Bücherl et al., 2017), the Golgi marker was GmMan1.2-mCherry (Nelson et al., 2007). The S-EPCS and V-EPCS markers were AtSYT1 (AT2G20990) and AtVAP27-1 (AT3G60600.1), respectively, cloned into pGWB228 as described in the plasmid cloning section. Constructs in *A. tumefaciens* were co-expressed at a ratio of protein of interest (POI):marker:p19 of 1:1:1 for EV-tagRFP, SP-mRFP-HDEL, REM1.2-mRFP, GmMAN1.2-mCherry and AtVAP27.1-tagRFP. AtSYT1-tagRFP had a lower signal and required a protein of interest (POI):marker:p19 ratio of 1:2:1.

Confocal images were captured with a Leica SP5 system equipped with an Argon and DPSS laser and hybrid detectors or a Leica STELLARIS 5 system equipped with a white light laser and hybrid detectors. For further details on excitation and emission wavelengths as well as tissue and downstream image processing please see the supplementary materials note 1.

### Cell death assay and trypan blue staining

The indicated constructs were transiently expressed in *N. benthamiana* via Agrobacterium mediated transformation. For cell death testing a ratio of NLR:AVR:p19 of 1:6:1 was used. Trypan blue staining to reveal tissue undergoing a cell death response was performed as previously described (Ma et al., 2011).

## Acknowledgements

We would like to thank Xiaoyou Hou from the Department of Plant and Microbial Biology at the University of Zürich for the *Arabidopsis thaliana* Col-0 cDNA sample. Imaging was performed with equipment maintained by the Center for Microscopy and Image Analysis, University of Zurich. This work was supported by grants 310030_ 204165 and 310030B_182833 from the Swiss National Science Foundation.

## Author contributions

J.I. and B.K. designed the research. J.I., L.K., S.F., and V.W. performed the experiments. J.I. and S.F. analyzed the data. J.I., L.K., and B.K. wrote the manuscript.

