## Supplementary Material for "The wheat NLR protein PM3b localizes to endoplasmic reticulum-plasma membrane contact sites and interacts with AVRPM3^b2/c2^ through its LRR domain"

### Supplementary note 1: Laser scanning confocal microscopy acquisition settings, plasmolysis conditions, and image processing

For LSCM with either the excitation wavelengths 458nm for CFP, 488nm for eGFP and 561nm for tagRFP, emissions were collected at 465-490nm for CFP, 495-545nm for eGFP and 575-625nm for tagRFP. Emission for each fluorophore was detected sequentially to avoid overlap of emission between channels. Time-lapse imaging was done in the XYT mode with 3-5 second intervals. Z-stacks were collected at the suggested system optimized thickness or at a set 2μm interval. For plasmolysis, leaf samples were incubated in a 4% NaCl hypotonic solution for 10-20 minutes on the glass slide before imaging.

All image processing was performed using FIJI (Fiji Is Just ImageJ) with the built-in standard tools. The look up table (LUT) used for pseudo-coloring of the images was done with the plugin color-blind-luts (Vellutini, B. C. (2015). Color blind friendly lookup tables for Fiji/ImageJ (v1.2.1). Zenodo. https://doi.org/10.5281/zenodo.7596751). For viewing and display of images in figures, contrast adjustments were made with the built-in enhance contrast function set to 0.35% saturated pixels and normalized. Background subtraction was occasionally performed equally in each fluorescence channel of the same image. Fluorescence intensity measurements were made on the raw TIFF files prior to contrast enhancements.

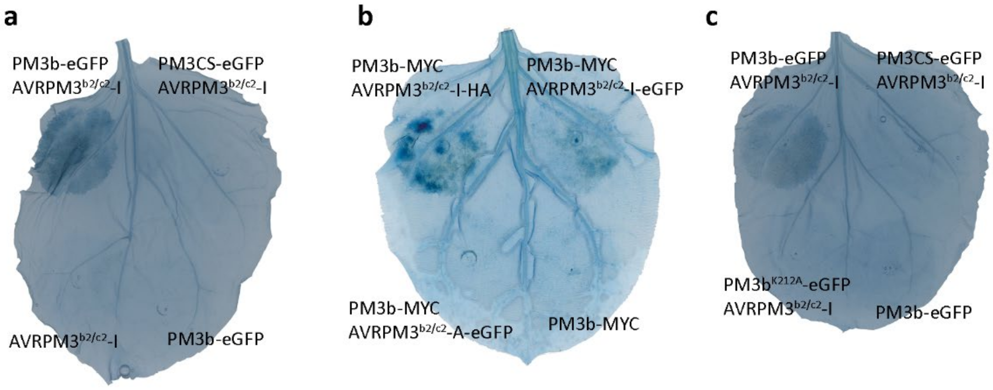


**Supplementary Fig. 1: Cell death induction capabilities of various PM3 variants and AVRPM3$^{b2/c2}$ variants used in this study. Trypan Blue staining was used to show cell death induction in N. benthamiana leafs 3-6 days post infiltration with Agrobacterium.** (a) AVRPM3$^{b2/c2}$ variants with different C-terminal fusions co-expressed with PM3b-MYC or PM3b-MYC alone as a negative control. (b) eGFP fused PM3b and PM3CS with variant I of AVRPM3$^{b2/c2}$ and AVRPM3$^{b2/c2}$ or PM3b-eGFP expressed alone as negative controls. (c) eGFP tagged PM3b variants co-expressed with AVRPM3$^{b2/c2}$-I showing abolished cell death induction of eGFP fused P-loop mutant of PM3b$^{K212A}$. PM3b-eGFP was used as a negative control.

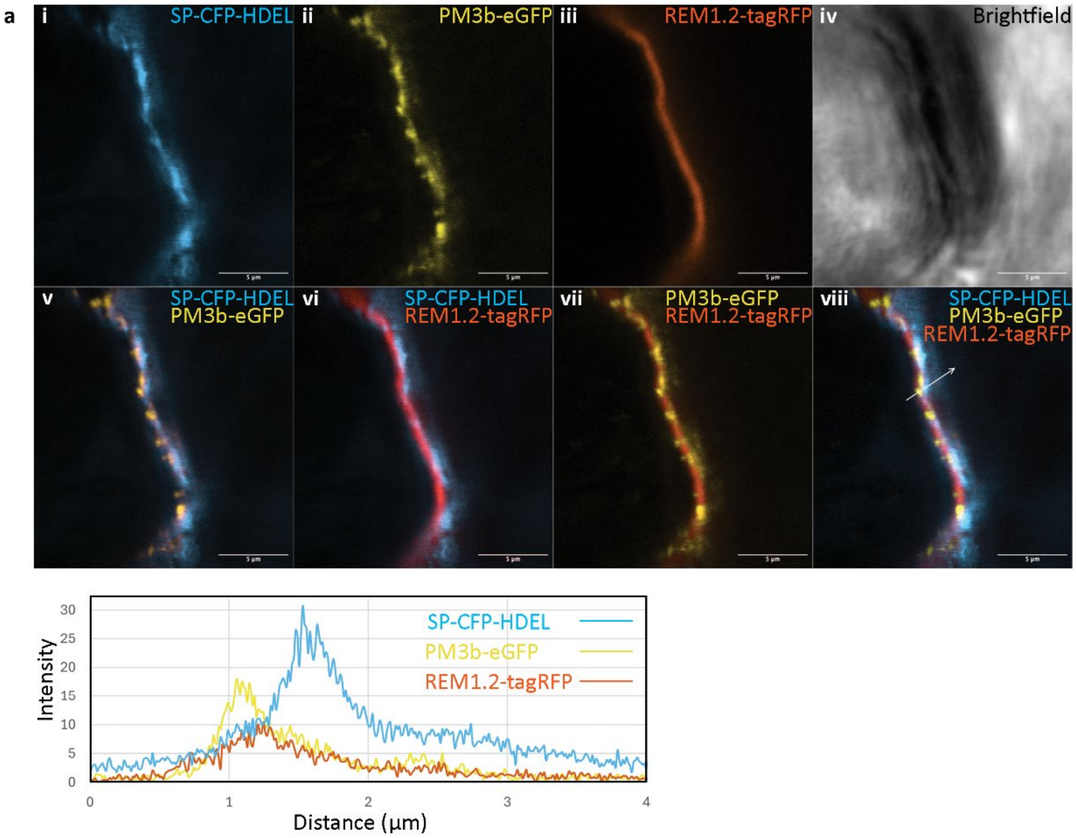


**Supplementary Fig. 2: PM3b puncta are partially overlapping with the plasma membrane and endoplasmic reticulum.** (a) Close up of the plasma membrane region of a cell expressing PM3b-eGFP, the ER marker SP-CFP-HDEL and the PM marker REM1.2-tagRFP. All combinations of merges between the acquired images of the fluorescence signals of each protein are shown. Fluorescence intensity histograms of the signals across a line of PM3b-eGFP, SP-CFP-HDEL and REM1.2-tagRFP were taken 48-72 hours post infiltration using laser scanning confocal microscopy. Each confocal image represents a single plane of a cell. Scale bars are 5µm.

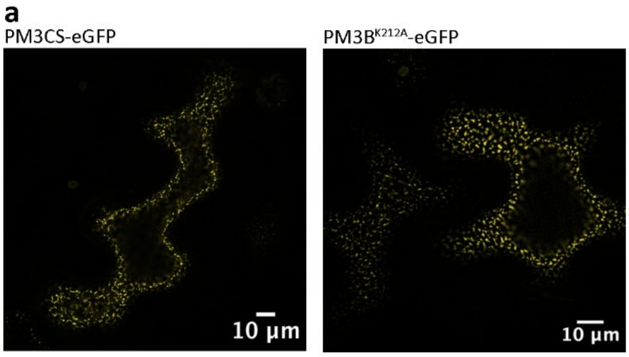


**Supplementary Fig. 3: PM3b$^{K212A}$ and PM3CS localize to puncta in N. benthamiana and wheat.** (a) eGFP tagged PM3b$^{K212A}$ and PM3CS were transiently expressed in N. benthamiana followed by laser scanning confocal microscopy at 48-72 hours post infiltration. The image is of a single plane of a cell. Scale bars are as indicated in each image.

**Supplementary table 1: DNA sequences of *Nlrs*, *Effectors* and *A. thaliana* genes cloned and used in this study.**

| Name | Sequence |
|---|---|
| NLR constructs | |
| Pm3b | ATGGCAGAGCGGGTGGTCACCATGGCGATCGGGCCACTGGTGTCCATGCTGAAGGACAAGGCG<br>TCCAGCTACCTCCTGGACCAGTACAAGGTCATGGAGGGAATGGAGGAGCAGCACAAGATTCTCA<br>AACGCAAGCTTCCGGCCATCCTCGACGTCATCACCGATGTCGAGGAGCAGGCCATGGCACAGAG<br>AGAAGGTGCCAAAGCCTGGCTTCAGGAGCTCAGGACAGTCGCCTATGTGGCAAATGAAGTCTTC<br>GACGAATTCAAGTACGAAGCCCTACGCCGTGAAGCCAAGAAGAATGGGCACTACATAAAGCTCG<br>GCTTCGATGTAATTAAACTCTTCCCTACTCACAACCGTGTTGCGTTCCGTTACAAAATGGGTCGCA<br>AGCTTTGCCTGATTCTGCAAGCCGTTGAGGTCCTCATAGCAGAGATGCAGGTCTTTGGGTTCAAG<br>TACCAACCACAGCCACCGGTGTCCAAAGAGTGGAGGCATACAGATTATGTTAGCATTGACCCACA<br>AGAAATTGCCAGCAGATCCAGACACGAAGATAAGAAGAACATTATTGGTATACTAGTTGATGAAG<br>CTAGCAATGCAGATCTCACAGTTGTTCCTGTGGTTGCAATGGGGGGCCTTGGCAAGACCACATTA<br>GCGCAACTCATATACAATGATCCTGAAATTCAGAAGCATTTCCAGTTGCTGCTCTGGGTTTGTGTCT<br>CTGATACCTTTGATGTGAACTCCCTGGCCAAGAGTATAGTTGAAGCATCCCCCAATAAGAATGTTG<br>ATACAGACAAACCACCATTGGCTAGACTTCAAAAACTGGTCAGCGGACAGCGGTATCTCCTTGTAT<br>TGGATGATGTTTGGGACAACAAAGAGTTACGTAAGTGGGAAAGGCTGAAGGTATGTCTTCAGCA<br>TGGTGGCATGGGCAGTGCAGTGTTGACAACAACTCGTGATAAACGAGTTGCTGAAATTATGGGT<br>GCAGATAGGGCAGCCTACAATCTCAATGCTTTGGAGGATCACTTCATAAAGGAAATTATTGTGGAT<br>AGAGCATTCAGTTCAGAGAATGGAAAGATTCCCGAGCTACTCGAGATGGTTGGTGAGATTGTGA<br>AGAGATGTTGTGGCTCTCCATTAGCTGCAAGTGCACTGGGCTCTGTACTTCGTACCAAGACCACC<br>GTGAAAGAATGGAATGCTATAGCATCTAGAAGCAGCATTTGCACTGAGGAAACTGGAATCTTGCC<br>AATACTCAAGCTTAGCTACAACGACTTGCCATCGCACATGAAGCAGTGCTTTGCTTTTTGTGCTGT<br>ATTTCCAAAGGATTACAAGATTGATGTAGCGAAGTTGATCCAACTATGGATCGCAAATGGCTTTAT<br>CCCTGAACACAAGGAAGATAGTCTTGAAACCATTGGACAACTTATTTTTGATGAGCTTGCTTCAA<br>GGTCATTCTTTCTGGATATAGAGAAGAGTAAAGAAGACTGGGAGTATTATTCCAGAACAACATGTA<br>AAATCCACGATCTTATGCATGATATTGCAATGTCTGTTATGGAAAAGGAATGTGTTGTTGCAACTAT<br>GGAACCAAGTGAAATTGAGTGGCTTCCAGATACTGCTCGGCATTTGTTCTTGTCATGTGAAGAAA<br>CGGAACGTATTTTGAATGATTCTATGGAGGAAAGATCCCCTGCTATTCAAACATTGCTATGTGATAG<br>TAATGTGTTCAGCCCATTGAAGCATCTATCAAAATATAGCTCTTTGCACGCCTTGAAGCTCTGTATTA<br>GAGGCACAGAATCATTTCTACTCAAACCAAAGTATCTGCATCACCTGAGGTACCTTGATCTCTCAG<br>AAAGTTCTATCAAAGCACTTCCTGAGGATATAAGTATTCTATATAACCTGCAAGTGTTGGACCTTTC<br>CTACTGTAATTATCTTGATCGACTTCCAAGGCAAATGAAGTATATGACTTCCCTCTGCCACCTCTACA<br>CTCATGGATGTCGGAACTTGAAGAGCATGCCTCCAGGACTTGAAAATCTCACTAAGCTGCAGACA<br>CTCACAGTTTTTGTAGCAGGAGTTCCTGGCCCTGATTGCGCTGATGTTGGAGAGCTGCATGGTCT<br>AAACATTGGTGGTCGGCTAGAGCTATGCCAGGTAGAGAATGTTGAAAAAGCAGAAGCAGAAGT<br>GGCAAACCTTGGTGGTCAGCTAGAGCTGCAGCATTTAAACCTTGGTGATCAGCTAGAGCTACGCC<br>GGGTAGAGAATGTTAAAAAAGCAGAGGCAAAAGTGGCGAATCTTGGAAACAAGAAGGATCTCC<br>GTGAACTGACATTAAGATGGACTGAGGTTGGCGACAGCAAGGTGCTCGACAAGTTCGAACCTCA<br>TGGTGGGCTGCAGGTTCTGAAGATATATAAATATGGAGGGAAGTGTATGGGTATGTTGCAAAACA<br>TGGTTGAGATCCATCTTTCTGGTTGTGAAAGATTGCAGGTTTTGTTCAGTTGTGGTACATCCTTCA<br>CGTTTCCAAAACTGAAGGTGCTTACACTAGAGCATTTATTGGATTTTGAGAGATGGTGGGAAATA<br>AATGAGGCACAAGAAGAACAGATAATATTTCCTCTGCTTGAGAAGTTGTTTATTCGGCATTGTGGA<br>AAGCTGATAGCATTACCTGAAGCACCATTGCTTGGAGAACCAAGTCGTGGAGGTAATAGACTGGT<br>ATGCACACCATTTTCTCTGCTTGAGAACTTGTTTATTTGGTATTGTGGAAAGCTGGTACCATTGCGT |

| | |
|---|---|
| | GAAGCACCACTGGTTCATGAAAGTTGTAGTGGAGGTTATAGGTTGGTACAGTCAGCATTTCCTGC<br>TCTAAAGGTACTTGCATTGGAAGACTTGGGGAGTTTTCAGAAATGGGATGCTGCTGTCGAAGGA<br>GAACCGATATTGTTTCCTCAGCTTGAGACACTATCAGTTCAGAAATGCCCAAAGCTAGTAGATTTA<br>CCCGAAGCACCAAAACTCAGTGTACTAGTAATTGAAGATGGCAAGCAAGAGGTGTTCCATTTTGT<br>AGACAGGTATTTATCTTCATTGACCAATCTGACACTGAGGCTAGAACACAGAGAAACAACATCAG<br>AGGCTGAATGCACTTCAATTGTACCTGTGGACAGCAAAGAGAAATGGAACCAGAAATCCCCTCTT<br>ACAGTTCTGGAGTTAGGATGCTGCAACTCATTCTTTGGACCAGGTGCACTAGAGCCGTGGGACTA<br>TTTTGTACACCTTGAAAAGTTGGAAATTGATAGATGTGATGTGCTCGTCCACTGGCCAGAGAATGT<br>GTTCCAAAGCTTGGTATCCTTGAGGACATTACTGATTAGAAACTGCAAAAATCTGACTGGATATGC<br>ACAAGCTCCTCTTGAGCCGTTGGCGTCTGAAAGGAGTCAGCACCCGAGAGGTCTGGAGTCTCTT<br>TGCTTAAGAAACTGCCCAAGTTTAGTAGAGATGTTCAACGTCCCGGCATCTCTCAAGAAAATGAC<br>TATTGGTGGGTGCATTAAGCTTGAGTCCATATTCGGCAAGCAACAGGGCATGGCAGAGTTAGTCC<br>AAGTATCTTCTAGCAGTGAGGCAATCATGCCTGCAACTGTATCAGAGTTGCCATCCACACCCATGA<br>ATCACTTTTGTCCATGCCTAGAAGATCTATGCTTATCAGCATGTGGAAGCTTACCAGCGGTTCTAAA<br>TCTGCCTCCATCCTTAAAGACCTTAGAAATGGATCGTTGTAGTAGTATTCAAGTCCTATCATGCCAG<br>CTGGGTGGGCTCCAGAAACCAGAAGCCACTACCTCCAGAAGCAGAAGTCCTATCATGCCACAGC<br>CACTAGCAGCAGCAACAGCACCAGCTGCAAGAGAGCATTTACTTCCTCCCCATCTCGAATATCTAA<br>CAATACTGAACTGTGCTGGCATGTTGGGTGGGACTCTCCGTCTGCCTGCACCCCTCAAGAGACTG<br>TTCATTATTGGCAACAGTGGGCTGACATCGCTGGAGTGTCTGTCGGGAGAGCACCCCCCATCGCT<br>GGAATCCCTTTGGCTTGAAAGATGCAGTACCCTGGCATCCCTGCCGAATGAGCCGCAAGTATACA<br>GGTCTCTCTGGTCTCTTGAAATTACAGGCTGCCCTGCTATAAAGAAGCTCCCTAGATGCCTGCAGC<br>AGCAACTGGGCAGCATCAAACGCAAATGGCTAGATGCCCGTTATGAAGTAACGGAATTCAAACCA<br>TTGAAACCGAAGACATGGAAGGAAATACCGAGGCTAGTCCGTGAGCGGAGGCAGGCCTGCCGG<br>AGC |
| Pm3b P-loop mutant K212A | ATGGCAGAGCGGGTGGTCACCATGGCGATCGGGCCACTGGTGTCCATGCTGAAGGACAAGGCG<br>TCCAGCTACCTCCTGGACCAGTACAAGGTCATGGAGGGAATGGAGGAGCAGCACAAGATTCTCA<br>AACGCAAGCTTCCGGCCATCCTCGACGTCATCACCGATGTCGAGGAGCAGGCCATGGCACAGAG<br>AGAAGGTGCCAAAGCCTGGCTTCAGGAGCTCAGGACAGTCGCCTATGTGGCAAATGAAGTCTTC<br>GACGAATTCAAGTACGAAGCCCTACGCCGTGAAGCCAAGAAGAATGGGCACTACATAAAGCTCG<br>GCTTCGATGTAATTAAACTCTTCCCTACTCACAACCGTGTTGCGTTCCGTTACAAAATGGGTCGCA<br>AGCTTTGCCTGATTCTGCAAGCCGTTGAGGTCCTCATAGCAGAGATGCAGGTCTTTGGGTTCAAG<br>TACCAACCACAGCCACCGGTGTCCAAAGAGTGGAGGCATACAGATTATGTTAGCATTGACCCACA<br>AGAAATTGCCAGCAGATCCAGACACGAAGATAAGAAGAACATTATTGGTATACTAGTTGATGAAG<br>CTAGCAATGCAGATCTCACAGTTGTTCCTGTGGTTGCAATGGGGGGCCTTGGCGCGACCACATTA<br>GCGCAACTCATATACAATGATCCTGAAATTCAGAAGCATTTCCAGTTGCTGCTCTGGGTTTGTGTCT<br>CTGATACCTTTGATGTGAACTCCCTGGCCAAGAGTATAGTTGAAGCATCCCCCAATAAGAATGTTG<br>ATACAGACAAACCACCATTGGCTAGACTTCAAAAACTGGTCAGCGGACAGCGGTATCTCCTTGTAT<br>TGGATGATGTTTGGGACAACAAAGAGTTACGTAAGTGGGAAAGGCTGAAGGTATGTCTTCAGCA<br>TGGTGGCATGGGCAGTGCAGTGTTGACAACAACTCGTGATAAACGAGTTGCTGAAATTATGGGT<br>GCAGATAGGGCAGCCTACAATCTCAATGCTTTGGAGGATCACTTCATAAAGGAAATTATTGTGGAT<br>AGAGCATTCAGTTCAGAGAATGGAAAGATTCCCGAGCTACTCGAGATGGTTGGTGAGATTGTGA<br>AGAGATGTTGTGGCTCTCCATTAGCTGCAAGTGCACTGGGCTCTGTACTTCGTACCAAGACCACC<br>GTGAAAGAATGGAATGCTATAGCATCTAGAAGCAGCATTTGCACTGAGGAAACTGGAATCTTGCC<br>AATACTCAAGCTTAGCTACAACGACTTGCCATCGCACATGAAGCAGTGCTTTGCTTTTTGTGCTGT<br>ATTTCCAAAGGATTACAAGATTGATGTAGCGAAGTTGATCCAACTATGGATCGCAAATGGCTTTAT<br>CCCTGAACACAAGGAAGATAGTCTTGAAACCATTGGACAACTTATTTTTGATGAGCTTGCTTCAA<br>GGTCATTCTTTCTGGATATAGAGAAGAGTAAAGAAGACTGGGAGTATTATTCCAGAACAACATGTA<br>AAATCCACGATCTTATGCATGATATTGCAATGTCTGTTATGGAAAAGGAATGTGTTGTTGCAACTAT |

| | |
|---|---|
| | GGAACCAAGTGAAATTGAGTGGCTTCCAGATACTGCTCGGCATTTGTTCTTGTCATGTGAAGAAA<br>CGGAACGTATTTTGAATGATTCTATGGAGGAAAGATCCCCTGCTATTCAAACATTGCTATGTGATAG<br>TAATGTGTTCAGCCCATTGAAGCATCTATCAAAATATAGCTCTTTGCACGCCTTGAAGCTCTGTATTA<br>GAGGCACAGAATCATTTCTACTCAAACCAAAGTATCTGCATCACCTGAGGTACCTTGATCTCTCAG<br>AAAGTTCTATCAAAGCACTTCCTGAGGATATAAGTATTCTATATAACCTGCAAGTGTTGGACCTTTC<br>CTACTGTAATTATCTTGATCGACTTCCAAGGCAAATGAAGTATATGACTTCCCTCTGCCACCTCTACA<br>CTCATGGATGTCGGAACTTGAAGAGCATGCCTCCAGGACTTGAAAATCTCACTAAGCTGCAGACA<br>CTCACAGTTTTTGTAGCAGGAGTTCCTGGCCCTGATTGCGCTGATGTTGGAGAGCTGCATGGTCT<br>AAACATTGGTGGTCGGCTAGAGCTATGCCAGGTAGAGAATGTTGAAAAAGCAGAAGCAGAAGT<br>GGCAAACCTTGGTGGTCAGCTAGAGCTGCAGCATTTAAACCTTGGTGATCAGCTAGAGCTACGCC<br>GGGTAGAGAATGTTAAAAAAGCAGAGGCAAAAGTGGCGAATCTTGGAAACAAGAAGGATCTCC<br>GTGAACTGACATTAAGATGGACTGAGGTTGGCGACAGCAAGGTGCTCGACAAGTTCGAACCTCA<br>TGGTGGGCTGCAGGTTCTGAAGATATATAAATATGGAGGGAAGTGTATGGGTATGTTGCAAAACA<br>TGGTTGAGATCCATCTTTCTGGTTGTGAAAGATTGCAGGTTTTGTTCAGTTGTGGTACATCCTTCA<br>CGTTTCCAAAACTGAAGGTGCTTACACTAGAGCATTTATTGGATTTTGAGAGATGGTGGGAAATA<br>AATGAGGCACAAGAAGAACAGATAATATTTCCTCTGCTTGAGAAGTTGTTTATTCGGCATTGTGGA<br>AAGCTGATAGCATTACCTGAAGCACCATTGCTTGGAGAACCAAGTCGTGGAGGTAATAGACTGGT<br>ATGCACACCATTTTCTCTGCTTGAGAACTTGTTTATTTGGTATTGTGGAAAGCTGGTACCATTGCGT<br>GAAGCACCACTGGTTCATGAAAGTTGTAGTGGAGGTTATAGGTTGGTACAGTCAGCATTTCCTGC<br>TCTAAAGGTACTTGCATTGGAAGACTTGGGGAGTTTTCAGAAATGGGATGCTGCTGTCGAAGGA<br>GAACCGATATTGTTTCCTCAGCTTGAGACACTATCAGTTCAGAAATGCCCAAAGCTAGTAGATTTA<br>CCCGAAGCACCAAAACTCAGTGTACTAGTAATTGAAGATGGCAAGCAAGAGGTGTTCCATTTTGT<br>AGACAGGTATTTATCTTCATTGACCAATCTGACACTGAGGCTAGAACACAGAGAAACAACATCAG<br>AGGCTGAATGCACTTCAATTGTACCTGTGGACAGCAAAGAGAAATGGAACCAGAAATCCCCTCTT<br>ACAGTTCTGGAGTTAGGATGCTGCAACTCATTCTTTGGACCAGGTGCACTAGAGCCGTGGGACTA<br>TTTTGTACACCTTGAAAAGTTGGAAATTGATAGATGTGATGTGCTCGTCCACTGGCCAGAGAATGT<br>GTTCCAAAGCTTGGTATCCTTGAGGACATTACTGATTAGAAACTGCAAAAATCTGACTGGATATGC<br>ACAAGCTCCTCTTGAGCCGTTGGCGTCTGAAAGGAGTCAGCACCCGAGAGGTCTGGAGTCTCTT<br>TGCTTAAGAAACTGCCCAAGTTTAGTAGAGATGTTCAACGTCCCGGCATCTCTCAAGAAAATGAC<br>TATTGGTGGGTGCATTAAGCTTGAGTCCATATTCGGCAAGCAACAGGGCATGGCAGAGTTAGTCC<br>AAGTATCTTCTAGCAGTGAGGCAATCATGCCTGCAACTGTATCAGAGTTGCCATCCACACCCATGA<br>ATCACTTTTGTCCATGCCTAGAAGATCTATGCTTATCAGCATGTGGAAGCTTACCAGCGGTTCTAAA<br>TCTGCCTCCATCCTTAAAGACCTTAGAAATGGATCGTTGTAGTAGTATTCAAGTCCTATCATGCCAG<br>CTGGGTGGGCTCCAGAAACCAGAAGCCACTACCTCCAGAAGCAGAAGTCCTATCATGCCACAGC<br>CACTAGCAGCAGCAACAGCACCAGCTGCAAGAGAGCATTTACTTCCTCCCCATCTCGAATATCTAA<br>CAATACTGAACTGTGCTGGCATGTTGGGTGGGACTCTCCGTCTGCCTGCACCCCTCAAGAGACTG<br>TTCATTATTGGCAACAGTGGGCTGACATCGCTGGAGTGTCTGTCGGGAGAGCACCCCCCATCGCT<br>GGAATCCCTTTGGCTTGAAAGATGCAGTACCCTGGCATCCCTGCCGAATGAGCCGCAAGTATACA<br>GGTCTCTCTGGTCTCTTGAAATTACAGGCTGCCCTGCTATAAAGAAGCTCCCTAGATGCCTGCAGC<br>AGCAACTGGGCAGCATCAAACGCAAATGGCTAGATGCCCGTTATGAAGTAACGGAATTCAAACCA<br>TTGAAACCGAAGACATGGAAGGAAATACCGAGGCTAGTCCGTGAGCGGAGGCAGGCCTGCCGG<br>AGC |
| Pm3b LRR domain alone | ATGTCTTTGCACGCCTTGAAGCTCTGTATTAGAGGCACAGAATCATTTCTACTCAAACCAAAGTATC<br>TGCATCACCTGAGGTACCTTGATCTCTCAGAAAGTTCTATCAAAGCACTTCCTGAGGATATAAGTAT<br>TCTATATAACCTGCAAGTGTTGGACCTTTCCTACTGTAATTATCTTGATCGACTTCCAAGGCAAATG<br>AAGTATATGACTTCCCTCTGCCACCTCTACACTCATGGATGTCGGAACTTGAAGAGCATGCCTCCA<br>GGACTTGAAAATCTCACTAAGCTGCAGACACTCACAGTTTTTGTAGCAGGAGTTCCTGGCCCTGA<br>TTGCGCTGATGTTGGAGAGCTGCATGGTCTAAACATTGGTGGTCGGCTAGAGCTATGCCAGGTAG |

| | |
|---|---|
| | AGAATGTTGAAAAAGCAGAAGCAGAAGTGGCAAACCTTGGTGGTCAGCTAGAGCTGCAGCATT<br>TAAACCTTGGTGATCAGCTAGAGCTACGCCGGGTAGAGAATGTTAAAAAAGCAGAGGCAAAAGT<br>GGCGAATCTTGGAAACAAGAAGGATCTCCGTGAACTGACATTAAGATGGACTGAGGTTGGCGAC<br>AGCAAGGTGCTCGACAAGTTCGAACCTCATGGTGGGCTGCAGGTTCTGAAGATATATAAATATGG<br>AGGGAAGTGTATGGGTATGTTGCAAAACATGGTTGAGATCCATCTTTCTGGTTGTGAAAGATTGC<br>AGGTTTTGTTCAGTTGTGGTACATCCTTCACGTTTCCAAAACTGAAGGTGCTTACACTAGAGCATT<br>TATTGGATTTTGAGAGATGGTGGGAAATAAATGAGGCACAAGAAGAACAGATAATATTTCCTCTG<br>CTTGAGAAGTTGTTTATTCGGCATTGTGGAAAGCTGATAGCATTACCTGAAGCACCATTGCTTGGA<br>GAACCAAGTCGTGGAGGTAATAGACTGGTATGCACACCATTTTCTCTGCTTGAGAACTTGTTTATT<br>TGGTATTGTGGAAAGCTGGTACCATTGCGTGAAGCACCACTGGTTCATGAAAGTTGTAGTGGAGG<br>TTATAGGTTGGTACAGTCAGCATTTCCTGCTCTAAAGGTACTTGCATTGGAAGACTTGGGGAGTTT<br>TCAGAAATGGGATGCTGCTGTCGAAGGAGAACCGATATTGTTTCCTCAGCTTGAGACACTATCAG<br>TTCAGAAATGCCCAAAGCTAGTAGATTTACCCGAAGCACCAAAACTCAGTGTACTAGTAATTGAA<br>GATGGCAAGCAAGAGGTGTTCCATTTTGTAGACAGGTATTTATCTTCATTGACCAATCTGACACTG<br>AGGCTAGAACACAGAGAAACAACATCAGAGGCTGAATGCACTTCAATTGTACCTGTGGACAGCA<br>AAGAGAAATGGAACCAGAAATCCCCTCTTACAGTTCTGGAGTTAGGATGCTGCAACTCATTCTTT<br>GGACCAGGTGCACTAGAGCCGTGGGACTATTTTGTACACCTTGAAAAGTTGGAAATTGATAGATG<br>TGATGTGCTCGTCCACTGGCCAGAGAATGTGTTCCAAAGCTTGGTATCCTTGAGGACATTACTGAT<br>TAGAAACTGCAAAAATCTGACTGGATATGCACAAGCTCCTCTTGAGCCGTTGGCGTCTGAAAGGA<br>GTCAGCACCCGAGAGGTCTGGAGTCTCTTTGCTTAAGAAACTGCCCAAGTTTAGTAGAGATGTTC<br>AACGTCCCGGCATCTCTCAAGAAAATGACTATTGGTGGGTGCATTAAGCTTGAGTCCATATTCGGC<br>AAGCAACAGGGCATGGCAGAGTTAGTCCAAGTATCTTCTAGCAGTGAGGCAATCATGCCTGCAAC<br>TGTATCAGAGTTGCCATCCACACCCATGAATCACTTTTGTCCATGCCTAGAAGATCTATGCTTATCA<br>GCATGTGGAAGCTTACCAGCGGTTCTAAATCTGCCTCCATCCTTAAAGACCTTAGAAATGGATCGT<br>TGTAGTAGTATTCAAGTCCTATCATGCCAGCTGGGTGGGCTCCAGAAACCAGAAGCCACTACCTCC<br>AGAAGCAGAAGTCCTATCATGCCACAGCCACTAGCAGCAGCAACAGCACCAGCTGCAAGAGAGC<br>ATTTACTTCCTCCCCATCTCGAATATCTAACAATACTGAACTGTGCTGGCATGTTGGGTGGGACTCT<br>CCGTCTGCCTGCACCCCTCAAGAGACTGTTCATTATTGGCAACAGTGGGCTGACATCGCTGGAGT<br>GTCTGTCGGGAGAGCACCCCCCATCGCTGGAATCCCTTTGGCTTGAAAGATGCAGTACCCTGGCA<br>TCCCTGCCGAATGAGCCGCAAGTATACAGGTCTCTCTGGTCTCTTGAAATTACAGGCTGCCCTGCT<br>ATAAAGAAGCTCCCTAGATGCCTGCAGCAGCAACTGGGCAGCATCAAACGCAAATGGCTAGATGC<br>CCGTTATGAAGTAACGGAATTCAAACCATTGAAACCGAAGACATGGAAGGAAATACCGAGGCTA<br>GTCCGTGAGCGGAGGCAGGCCTGCCGGAGC |
| Pm3CS | ATGGCAGAGCGGGTGGTCACCATGGCGATCGGGCCACTGGTGTCCATGCTGAAGGACAAGGCG<br>TCCAGCTACCTCCTGGACCAGTACAAGGTCATGGAGGGAATGGAGGAGCAGCACAAGATTCTCA<br>AACGCAAGCTTCCGGCCATCCTCGACGTCATCACCGATGTCGAGGAGCAGGCCATGGCACAGAG<br>AGAAGGTGCCAAAGCCTGGCTTCAGGAGCTCAGGACAGTCGCCTATGTGGCAAATGAAGTCTTC<br>GACGAATTCAAGTACGAAGCCCTACGCCGTGAAGCCAAGAAGAATGGGCACTACATAAAGCTCG<br>GCTTCGATGTAATTAAACTCTTCCCTACTCACAACCGTGTTGCGTTCCGTTACAAAATGGGTCGCA<br>AGCTTTGCCTGATTCTGCAAGCCGTTGAGGTCCTCATAGCAGAGATGCAGGTCTTTGGGTTCAAG<br>TACCAACCACAGCCACCGGTGTCCAAAGAGTGGAGGCATACAGATTATGTTAGCATTGACCCACA<br>AGAAATTGCCAGCAGATCCAGACACGAAGATAAGAAGAACATTATTGGTATACTAGTTGATGAAG<br>CTAGCAATGCAGATCTCACAGTTGTTCCTGTGGTTGCAATGGGGGGCCTTGGCAAGACCACATTA<br>GCGCAACTCATATACAATGATCCTGAAATTCAGAAGCATTTCCAGTTGCTGCTCTGGGTTTGTGTCT<br>CTGATACCTTTGATGTGAACTCCCTGGCCAAGAGTATAGTTGAAGCATCCCCCAATAAGAATGTTG<br>ATACAGACAAACCACCATTGGATAGACTTCAAAAACTGGTCAGCGGACAGCGGTATCTCCTTGTAT<br>TGGATGATGTTTGGGACAACAAAGAGTTACGTAAGTGGGAAAGGCTGAAGGTATGTCTCCAGCA<br>TGGTGGCATGGGCAGTGCAGTGTTGACAACAACTCGTGATAAACGAGTTTCTGAAATTATGGGTG |

| | |
|---|---|
| | CAGATAGAGCTGCCTACAATCTCAATGCTTTGGAGGATCACTTCATAAAGGAAATTATTGAGGCTA<br>GAGCGTTCAGTTCAAAGAAAGAAAAGCCTATTGAACTAGTGGAGGTAGTTGACGAGATTGTGAA<br>GAGATGTTGTGGCTCTCCATTAGCTGCAACGGCACTGGGCTCTGTACTTTGTACCAAGACCAGTG<br>TGAAAGAATGGAAAGCTGTATCATCTGGAACCAGCGTTTGCACTGATGAAACTGGAATTTTGCCA<br>ATACTCAAGCTTAGCTACAACGACTTGCCCGCACACATGAAGCAGTGCTTTGCTTTCTGTGCTGTG<br>TTTCCCAAGGATTACAAGATCAATGTGGAGAAGCTTATCCAACTATGGATTGCAAACGGCTTTATC<br>CTAGAATACAAGGAAGATAGTCCCGAAACATTTGGAAAACATATTTTCGATGAGCTTGTCTCAAGG<br>TCATTCTTTCTGGACCTAGAGGAAAGTAAGGACTACAGCGGGTATTATTCCAGCACATGTAAAATT<br>CATGATCTTATGCATGATATTGCAATGTCTGTTATGGAAAAGGAATGTGTTGTTGCAACTATGGAAC<br>CAAGTGAAATTGAGTGGCTTCCAGATACTGCTCGGCATTTGTTCTTGTCATGTGAAGAAGCAGAA<br>CGTATTTTGAATGATTCTATGCAGGAAAGATCCCCTGCTATTCAAACATTGCTATGCAATAGTGATG<br>TGTTCAGCCCATTGCAGCATCTATCAAAATACAACACTTTGCATGCCTTGAAGCTCTGTCTGGGAA<br>CAGAATCATTTCTACTCAAACCAAAGTATCTGCATCACCTGAGGTACCTTGATCTCTCAGAAAGTTC<br>TATCAAAGCACTTCCTGAGGATATAAGTATTCTATATAACCTGCAAGTGTTGGACCTTTCCTACTGTA<br>ATTATCTTGATCGACTTCCAAGGCAAATGAAGTATATGACTTCCCTCTGCCACCTCTACACTCATGG<br>ATGTCGGAACTTGAAGAGCATGCCTCCAGGACTTGAAAATCTCACTAAGCTGCAGACACTCACAG<br>TTTTTGTAGCAGGAGTTCCTGGCCCTGATTGCGCTGATGTTGGAGAGCTGCATGGTCTAAACATT<br>GGTGGTCGGCTAGAGCTATGCCAGGTAGAGAATGTTGAAAAAGCAGAAGCAGAAGTGGCAAAC<br>CTTGGTGGTCAGCTAGAGCTGCAGCATTTAAACCTTGGTGATCAGCTAGAGCTACGCCGGGTAGA<br>GAATGTTAAAAAAGCAGAGGCAAAAGTGGCGAATCTTGGAAACAAGAAGGATCTCCGTGAACT<br>GACATTAAGATGGACTGAGGTTGGCGACAGCAAGGTGCTCGACAAGTTCGAACCTCATGGTGGG<br>CTGCAGGTTCTGAAGATATATAAATATGGAGGGAAGTGTATGGGTATGTTGCAAAACATGGTTGA<br>GATCCATCTTTCTGGTTGTGAAAGATTGCAGGTTTTGTTCAGTTGTGGTACATCCTTCACGTTTCCA<br>AAACTGAAGGTGCTTACACTAGAGCATTTATTGGATTTTGAGAGATGGTGGGAAATAAATGAGGC<br>ACAAGAAGAACAGATAATATTTCCTCTGCTTGAGAAGTTGTTTATTCGGCATTGTGGAAAGCTGAT<br>AGCATTACCTGAAGCACCATTGCTTGGAGAACCAAGTCGTGGAGGTAATAGACTGGTATGCACAC<br>CATTTTCTCTGCTTGAGAACTTGTTTATTTGGTATTGTGGAAAGCTGGTACCATTGCGTGAAGCAC<br>CACTGGTTCATGAAAGTTGTAGTGGAGGTTATAGGTTGGTACAGTCAGCATTTCCTGCTCTAAAG<br>GTACTTGCATTGGAAGACTTGGGGAGTTTTCAGAAATGGGATGCTGCTGTCGAAGGAGAACCGA<br>TATTGTTTCCTCAGCTTGAGACACTATCAGTTCAGAAATGCCCAAAGCTAGTAGATTTACCCGAAG<br>CACCAAAACTCAGTGTACTAGTAATTGAAGATGGCAAGCAAGAGGTGTTCCATTTTGTAGACAGG<br>TATTTATCTTCATTGACCAATCTGACACTGAGGCTAGAACACAGAGAAACAACATCAGAGGCTGA<br>ATGCACTTCAATTGTACCTGTGGACAGCAAAGAGAAATGGAACCAGAAATCCCCTCTTACAGTTC<br>TGGAGTTAGGATGCTGCAACTCATTCTTTGGACCAGGTGCACTAGAGCCGTGGGACTATTTTGTA<br>CACCTTGAAAAGTTGGAAATTGATAGATGTGATGTGCTCGTCCACTGGCCAGAGAATGTGTTCCA<br>AAGCTTGGTATCCTTGAGGACATTACTGATTAGAAACTGCAAAAATCTGACTGGATATGCACAAGC<br>TCCTCTTGAGCCGTTGGCGTCTGAAAGGAGTCAGCACCCGAGAGGTCTGGAGTCTCTTTGCTTAA<br>GAAACTGCCCAAGTTTAGTAGAGATGTTCAACGTCCCGGCATCTCTCAAGAAAATGACTATTGGT<br>GGGTGCATTAAGCTTGAGTCCATATTCGGCAAGCAACAGGGCATGGCAGAGTTAGTCCAAGTATC<br>TTCTAGCAGTGAGGCAATCATGCCTGCAACTGTATCAGAGTTGCCATCCACACCCATGAATCACTT<br>TTGTCCATGCCTAGAAGATCTATGCTTATCAGCATGTGGAAGCTTACCAGCGGTTCTAAATCTGCCT<br>CCATCCTTAAAGACCTTAGAAATGGATCGTTGTAGTAGTATTCAAGTCCTATCATGCCAGCTGGGTG<br>GGCTCCAGAAACCAGAAGCCACTACCTCCAGAAGCAGAAGTCCTATCATGCCACAGCCACTAGCA<br>GCAGCAACAGCACCAGCTGCAAGAGAGCATTTACTTCCTCCCCATCTCGAATATCTAACAATACTG<br>AACTGTGCTGGCATGTTGGGTGGGACTCTCCGTCTGCCTGCACCCCTCAAGAGACTGTTCATTATG<br>GGCAACAGTGGGCTGACATCGCTGGAGTGTCTGTCGGGAGAGCACCCCCCATCGCTGGAATCCC<br>TTTGGCTTGAAAGATGCAGTACCCTGGCATCCCTGCCGAATGAGCCGCAAGTATACAGGTCTCTCT<br>GGTCTCTTGAAATTACAGGCTGCCCTGCTATAAAGAAGCTCCCTAGATGCCTGCAGCAGCAACTG |

| | |
|---|---|
| | GGCAGCATCAAACGCAAATGGCTAGATGCCCGTTATGAAGTAACGGAATTCAAACCATTGAAACC<br>GAAGACATGGAAGGAAATACCGAGGCTAGTCCGTGAGCGGAGGCAGGCCTGCCGGAGC |
| Mla1 | ATGGATATTGTCACCGGTGCCATTTCCAACCTGATTCCCAAGTTGGGGGAGCTGCTCACGGAGGA<br>GTTCAAGCTGCACAAGGGTGTCAAGAAAAATATTGAGGACCTCGGGAAGGAGCTTGAGAGCAT<br>GAACGCTGCCCTCATCAAGATTGGTGAGGTGCCGAGGGAGCAGCTCGACAGCCAAGACAAGCT<br>CTGGGCCGATGAAGTCAGAGAGCTCTCCTACGTCATTGAGGATGTCGTCGACAAGTTCCTCGTAC<br>AGGTTGATGGCATTCAGTTTGATGATAACAACAACAAATTTAAGGGGTTCATGAAGAGGACGACC<br>GAGTTGTTGAAGAAAGTCAAGCATAAGCATGGGATAGCTCACGCGATCAAGGACATCCAAGAGC<br>AACTCCAAAAGGTGGCTGATAGGCGTGACAGGAACAAGGTATTTGTTCCTCATCCTACGAGAACA<br>ATTGCTATTGACCCTTGCCTTCGAGCTTTGTATGCTGAAGCGACAGAGCTAGTTGGCATATATGGA<br>AAGAGGGATCAAGACCTCATGAGGTTGCTTTCCATGGAGGGCGATGATGCCTCTAATAAGAGACT<br>GAAGAAGGTCTCCATTGTTGGATTTGGAGGGTTGGGCAAGACCACTCTTGCTAGAGCGGTATAC<br>GAGAAGATTAAAGGTGATTTCGATTGTCGGGCTTTTGTTCCGGTCGGTCAGAACCCTCACATGAA<br>GAAGGTTTTAAGGGATATCCTCATTGATCTCGGAAATCCTCACTCAGATCTTGCGATGCTGGATGC<br>CAATCAGCTTATTAAAAAACTTCGTGAATTTCTAGAGAACAAAAGGTATCTTGTCATAATTGATGAT<br>ATATGGGATGAAAAATTATGGGAAGGCATCAACTTTGCTTTCTCCAATAGGAATAATCTAGGCAGT<br>CGGCTAATCACCACAACCCGCATTGTCAGTGTCTCTAATTCATGTTGCTCATCACATGGTGATTCGG<br>TTTATCAAATGGAACCACTTTCTGTTGATGACTCCAGAATACTCTTCTGGAAAAGAATATTTCCAGA<br>TGAGAATGGATGTCTAAATGAATTTGAACAAGTGTCGAGAGATATTCTAAAGAAATGTGGTGGGG<br>TACCACTAGCCATAATTACCATAGCTAGTGCTTTGGCCGGTGACCAGAAGATGAAACCAAAGTGTG<br>AGTGGGATATTCTCCTTCAGTCCCTTGGCTCTGGACTAACAGAAGATAACAGTTTAGAGGAGATG<br>CGGAGAATACTCTCTTTCAGCTATTCTAATCTACCTTCTCATCTGAAAACTTGTCTACTGTATCTATGT<br>ATATATCCAGAAGATAGCAAGATTCATAGAGATGAACTGATATGGAAGTGGGTGGCCGAAGGATT<br>TGTCCACCATGAAAACCAAGGAAATAGCTTGTATTTGCTCGGATTAAATTACTTCAACCAGCTCATT<br>AATAGAAGTATGATCCAGCCCATATATGGTTTTAATGACGAGGTATATGTATGTCGTGTACATGATAT<br>GGTTCTGGACCTTATCTGCAACTTGTCACGTGAAGCAAAATTTGTGAATCTATTGGATGGCAGTGG<br>GAATAGCATGTCTTCACAGGGTAATTGTCGCCGTCTGTCCCTTCAAAAAAGAAATGAAGATCATCA<br>AGCCAAACCTATCACAGATATCAAGAGTATGTCACGAGTGAGGTCAATTACTATCTTTCCACCTGCT<br>ATTGAAGTCATGCCATCTCTTTCAAGGTTTGACGTTTTACGTGTACTTGATCTGTCACGATGTAATC<br>TTGGGGAGAATAGCAGCCTGCAGCTTAACCTGAAGGATGTTGGACATTTAACTCACCTAAGGTAC<br>CTTGGTCTAGAAGGTACCAACATCAGTAAGCTCCCTGCTGAGATAGGAAAACTGCAGTTTTTGGA<br>GGTGTTGGATCTTGGAAACAATCATAATCTAAAGGAATTGCCGTCCACTGTTTGTAATTTCAGAAG<br>ATTAATCTACCTAAATTTATTTGGGTGTCCGGTGGTTCCTCCAGTTGGTGTGTTGCAAAATCTGACA<br>TCCATAGAAGTGTTGAGGGGGATCTTGGTCTCTGTGAACATTATTGCACAAGAGCTTGGCAACCT<br>GGAAAGGCTGAGGGTGCTTGATATTTGCTTCAGGGATGGTAGTTTGGATTTGTATAAAGATTTCG<br>TGAAGTCTCTGTGCAACCTACATCACATCGAAAGTCTACGTATTGAGTGCAATTCCAGAGAAACAT<br>CATCTTTTGAACTGGTGGATCTCTTGGGAGAACGCTGGGTGCCTCCTGTACATTTCCGTGAATTTG<br>TGTCATCCATGCCTAGCCAACTCTCTGCACTGCGAGGGTGGATAAAGAGAGACCCCTCCCATCTCT<br>CGAACCTCTCCGAGTTAATCCTCTCGTCAGTGAAGGACGTGCAGCAGGATGACGTGGAAATCATT<br>GGGGGGTTGTTGTGCCTTCGTCGTCTCTTTATAATAACGAGCACCGACCAAACGCAACGGCTGCT<br>AGTCATCCGTGCAGATGGGTTCCGCTGTACGGTTGACTTTCGATTGGATTGTGGATCTGCCACGCA<br>GATATTGTTTGAACCAGGAGCTTTGCCAAGGGCGGTAAGAGTTTGGTTCAGCCTTGGCGTGCGG<br>GTGACGAAAGAGGATGGTAACCGTGGCTTCGACTTGGGCCTGCAGGGGAACCTGTTCTCCCTTC<br>GAGAGTTTGTCTCTGTTTATATGTATTGTGGTGGAGCGAGGGTTGGGGAGGCAAAGGAAGCGGA<br>GGCTGCGGTGAGGCGTGCCCTGGAAGCTCATCCCAGCCATCCCCGGATTTATATTCAGATGAGGC<br>CGCATATAGCAAAAGGTGCTCATGATGACGATTTGTGTGAGGACGAGGAGGAGAACGAC |
| Effector constructs | |

| | |
|---|---|
| AvrPm3b2/<br>c2_A | ATGTATTTGTTTTACCGATGCGGCAATGATTATATCACAGAGCGAGCTTTAATCAACCAGATTAGCA<br>TGGAACATAAGAAGTTAACCGGACAGGGCTCTAGTGCTGATTCCTTCCCTGGCGGGCGTGCCACT<br>GCCGAAGTTACGTTTTGGGAGCCCAGTATTTCTAACCCTGGGACTTATCTCGACATAAAAGTGAA<br>GTTTGACATATACAGGCAGATGCTGAGCTTTGAAGTATCCAGTTCTGGTAAGCGAATACCGTGTGA<br>AGGAGATTATGGCGCCGAGATACCTGAGGAAGACTTGGAAGTGTCAGACGAGCCTTATTATGCTA<br>ACTAG |
| AvrPm3b2/<br>c2_C | ATGTACTTGTTTTATCGATGTGGTAATGATTATATCACAGAAAGAGCTTTAATAGATCAAATTTCAAT<br>GGAGCACAAAAAACTCACAGGGCAAGGGAGCAGCGCAGACTCTTTTCCGGGAGGTCGAGCCA<br>CTGCTGAGGTAACATTCTGGGAGCCATCAATATCAAATCCCGGCACATATCTGGATATAAAGGTTAA<br>ATTCGACATATACAGACAAATGCTCAGCTTTGAGGTAAGTTCTAGCGGCAAGAGGATTCCGTGCG<br>AAGGCGACTACGGAGCCGAGATCCCGGAGGAAGATCTCGAAGTGTCAGACGAGCCCTACTACGC<br>CAATTAG |
| AvrPm3b2/<br>c2_I | ATGTACTTATTTTACAGATGTGGTAATGATTATATTACCGAAAGGGCACTGATAAATCAGATTAACAT<br>GGAACACAAGAAGTTGACGGGACAGGGCTCTTCCGCTGACAGTTTCCCTGGAGGGAGAGCTAC<br>AGCCGAAGTCACGTTTTGGGAGCCTTCAATAAGCAATCCTGGAACATACCTCGACATTAAAGTCA<br>AATTCGACATTTACCGACAAATGCTTTCTTTCGAGGTGTCAAGTAGTGGAAAGAGGATACCCTGT<br>GAAGGGGATTATGGAGCAGAAATCCCGGAAGAGGACTTGGAGGTCTCCGATGAGCCATATTATG<br>CCAACTAA |
| *Arabidopsis thaliana* ER-PM contact site marker constructs | |
| AtSyt1 | ATGGGCTTTTTCAGTACGATACTAGGATTTTGTGGATTTGGAGTTGGGATTTCATTGGGACTTGTT<br>ATTGGTTACGTTCTCTTCGTCTACTTGCTCCCCAACGACGTCAAGGATCCTGAAATTCGTTCAATAG<br>CTGACCAAGATCCCAAAGCTATGCTACGGATGCTTCCAGAGATACCTCTATGGGTCAAAAATCCGG<br>ATTTCGATCGTGTTGACTGGATAAACAGATTTCTCGAGTACATGTGGCCTTATCTCGACAAGGCCA<br>TATGTAAGACTGCAAAGAATATAGCAAAACCGATCATTGAGGAGCAGATACCAAAGTACAAGATT<br>GACTCTGTTGAATTTGAAACACTTACTCTAGGCTCTTTACCCCCTACATTTCAAGGGATGAAAGTTT<br>ATCTCACGGATGAGAAAGAGTTGATTATGGAACCATGCTTGAAATGGGCCGCAAATCCCAATATCT<br>TGGTTGCCATCAAGGCATTTGGGTTGAAGGCAACAGTTCAGGTGGTCGATCTGCAAGTTTTTGCT<br>CAGCCTCGTATCACTCTCAAGCCATTAGTTCCAAGTTTCCCTTGTTTTGCCAATATCTATGTGTCTCT<br>TATGGAGAAGCCACATGTTGATTTTGGACTGAAGCTTGGTGGAGCAGATCTTATGTCAATCCCTG<br>GCCTCTACAGATTTGTTCAGGAGCAAATCAAGGATCAAGTTGCGAACATGTATCTCTGGCCTAAGA<br>CCCTTGTAGTTCCAATCCTTGACCCTGCAAAGGCGTTCAGAAGGCCTGTTGGAATTGTCCATGTGA<br>AAGTTGTGAGGGCTGTGGGGCTGAGGAAAAAAGATCTGATGGGCGGGGCAGATCCATTCGTGA<br>AAATCAAGCTCTCTGAAGATAAGATTCCTTCTAAGAAGACAACAGTTAAACACAAGAATTTGAATC<br>CTGAATGGAATGAGGAGTTCAAATTCTCGGTCAGAGATCCCCAGACTCAGGTTCTAGAGTTCAGT<br>GTGTATGACTGGGAACAGGTTGGGAATCCCGAGAAGATGGGTATGAATGTATTAGCTCTGAAAGA<br>AATGGTGCCTGACGAACATAAAGCATTCACCTTGGAACTGCGTAAGACTCTGGACGGCGGTGAA<br>GATGGGCAGCCTCCTGACAAGTATAGGGGGAAGCTGGAGGTTGAACTCTTGTATAAGCCATTCAC<br>GGAGGAAGAAATGCCCAAAGGCTTTGAAGAAACGCAAGCCGTGCAGAAAGCTCCAGAAGGCA<br>CACCGGCTGCTGGAGGAATGCTTGTGGTAATAGTGCATTCGGCTGAGGATGTTGAAGGAAAGCA<br>CCATACCAATCCTTACGTGCGCATCTATTTCAAAGGAGAAGAGAGAAAAACAAAGCACGTGAAGA<br>AGAACAGAGACCCAAGGTGGAATGAGGAGTTCACCTTTATGCTCGAGGAGCCTCCAGTCCGTGA<br>GAAGCTGCACGTTGAAGTGCTGAGCACCTCTTCCAGGATAGGTCTATTGCATCCCAAGGAAACAC<br>TGGGGTATGTGGATATTCCAGTGGTGGACGTGGTGAACAACAAAAGGATGAATCAGAAGTTTCA<br>CCTTATTGATTCTAAGAACGGAAAGATCCAAATCGAGCTCGAGTGGCGAACTGCCTCT |
| AtVap27-1 | ATGAGTAACATCGATCTGATTGGGATGAGTAACCGCGATCTGATCGGGATGAGTAACAGCGAGCT<br>TCTCACCGTCGAGCCTCTCGATCTTCAATTCCCTTTTGAATTGAAGAAGCAGATCTCTTGCTCTCTC |

| | |
|---|---|
| | TATTTGACGAACAAGACCGACAATAATGTTGCCTTTAAGGTTAAGACGACGAATCCGAAAAAGTA<br>TTGTGTTAGGCCTAATACTGGAGTTGTTCTCCCGAGGTCTACTTGCGAAGTTCTTGTGACCATGCA<br>AGCTCAAAAGGAAGCTCCTTCCGATATGCAGTGCAAGGACAAGTTTCTGCTTCAAGGTGTGATAG<br>CTAGTCCTGGTGTCACAGCCAAGGAAGTTACTCCTGAGATGTTTAGCAAAGAGGCTGGACATCGA<br>GTTGAGGAAACTAAACTGAGAGTTACTTATGTTGCTCCACCACGACCACCATCACCGGTTCACGA<br>AGGATCTGAAGAGGGTTCTTCACCCAGGGCTTCTGTCTCAGATAATGGACATGGTTCTGAATTTTC<br>GTTTGAGAGATTTATCGTGGACAACAAGGCTGGACATCAAGAAAACACATCTGAGGCGAGGGCA<br>CTCATTACCAAGCTAACCGAAGAAAAACAGTCTGCCATTCAACTGAACAACAGACTTCAAAGAGA<br>ACTGGATCAGTTAAGGCGTGAAAGCAAAAAGAGCCAAAGTGGTGGTATCCCATTCATGTACGTTC<br>TTTTGGTCGGACTCATCGGTCTAATTTTGGGATACATTATGAAGAGGACA |
